# Lysosomal uptake of mtDNA mitigates heteroplasmy

**DOI:** 10.1101/2024.02.16.580263

**Authors:** Parisa Kakanj, Mari Bonse, Aylin Gökmen, Felix Gaedke, Belén Mollá, Elisabeth Vogelsang, Astrid Schauss, Andreas Wodarz, David Pla-Martín

## Abstract

Mitochondrial DNA is exposed to multiple insults produced by normal cellular function. Upon mtDNA replication stress the mitochondrial genome transfers to endosomes where it is degraded. Here, using proximity proteomics we found that mtDNA replication stress leads to the rewiring of the mitochondrial proximity proteome, increasing mitochondria association with lysosomal and vesicle-associated proteins, such as the GTPase RAB10 and the retromer. We found that upon mtDNA replication stress, RAB10 enhances mitochondrial fragmentation and relocates from the ER to lysosomes containing mtDNA. The retromer enhances and coordinates the expulsion of mitochondrial matrix components through mitochondrial-derived vesicles, and mtDNA with direct transfer to lysosomes. Using a *Drosophila* model carrying a long deletion on the mtDNA (ΔmtDNA), we evaluated *in vivo* the role of the retromer in mtDNA extraction and turnover in the larval epidermis. The presence of ΔmtDNA elicits the activation of a specific transcriptome profile related to counteract mitochondrial damage. Expression of the retromer component *Vps35* is sufficient to restore mtDNA homoplasmy and mitochondrial defects associated with ΔmtDNA. Our data reveal novel regulators involved in the specific elimination of mtDNA. We demonstrate that modulation of the retromer *in vivo* is a successful mechanism to restore mitochondrial function associated with mtDNA damage.

## Introduction

The presence of the mitochondrial genome (mtDNA) outside the mitochondrial matrix serves as a primary sign of mitochondrial dysfunction. The mtDNA, due to its similarity to the bacterial genome, can be recognized by intracellular defense systems^1^. Thus, rapid elimination of damaged mtDNA is crucial to avoid the engagement of cytoplasmic DNA sensors, which in turn, activate the innate immune response.

Several sources of mitochondrial dysfunction trigger mtDNA leakage. The accumulation of oxidative damage in the mtDNA modifies the biochemical properties of the molecule, alters base-pairing properties, and affects the maintenance of the mitochondrial genome, serving as a source for mutations^2, 3^. Thus, oxidative stress, produced either by normal cellular function or by external sources promotes mtDNA leak^4^. Upon metabolic stress, mtDNA escapes through mitochondrial-derived vesicles (MDVs) which are integrated into the cellular vesicular network^5^. In addition, interfering with mtDNA replication leads to the translocation of mitochondrial nucleoids to endosomes which are finally degraded into multivesicular bodies^6, 7^.

The presence of mtDNA in autophagy-related structures suggests a specific selective quality control pathway. Indeed, the proteomic profile of autophagy organelles in neurons suggests that mitochondrial nucleoids are selected for basal autophagy^8^. Although little is known about specific regulators to remove the mitochondrial genome upon mtDNA damage, recent studies indicate the involvement of vesicle trafficking and mitochondrial membrane components. Thus, RAB5, and in particular RAB5C, seems to participate in cargo selection. RAB5C physically interacts with the mitochondrial membrane proteins Mitofusin 1 and Mitofusin 2 and assists in endosomal approach and mtDNA transfer^9^. Similarly, upon mtDNA replication stress, mtDNA delivery to VPS35-RAB5 endosomes requires the mitochondrial membrane protein SAMM50^6^.

In the current study, we sought to elucidate the mechanisms of mtDNA elimination. Here we report that RAB10 and the retromer facilitate mitochondrial fragmentation and lysosomal recruitment upon mtDNA replication stress. Mitochondrial matrix components exit the mitochondrial network through mitochondrial-derived vesicles which are rapidly integrated in the lysosomes. mtDNA however transfers directly into lysosomes. This process is controlled and enhanced by the retromer component VPS35 and inhibited in cells expressing the Parkinson’s-associated mutation VPS35^D620N^. Further, using a Drosophila model containing a long deletion in the mtDNA (ΔmtDNA), we validate the role of *Vps35* in reducing mtDNA heteroplasmy. Our results reveal novel regulators for mtDNA turnover, highlighting the lysosomes as the central organelles maintaining mitochondrial function.

## Results

### mtDNA damage changes the proteome of mitochondria-endosome contact sites

Specific removal of mtDNA requires the recruitment of endosomal vesicles to the mitochondrial surface^6^. To detect novel regulators involved in mtDNA turnover, we used the promiscuous biotinylation enzyme Turbo ID^10^. For specific detection of the mitochondria-endosome proteome, we fused the N-terminal part of Turbo ID to RAB5C (TurboID aa1-aa72; RAB5C-SplitTurboNt-V5, Figure 1A) and, the C-terminal part to SAMM50 (TurboID aa73–aa246; SAMM50-SplitTurboCt-HA; Figure 1A). The expected subcellular localization of both transgene-encoded fusion proteins at endosomes and mitochondria respectively, was validated in HeLa cells with confocal microscopy (Fig. S1A and B). Reconstitution of biotinylation activity was also confirmed by western blot analysis (Fig. S1C).

**Figure 1.**
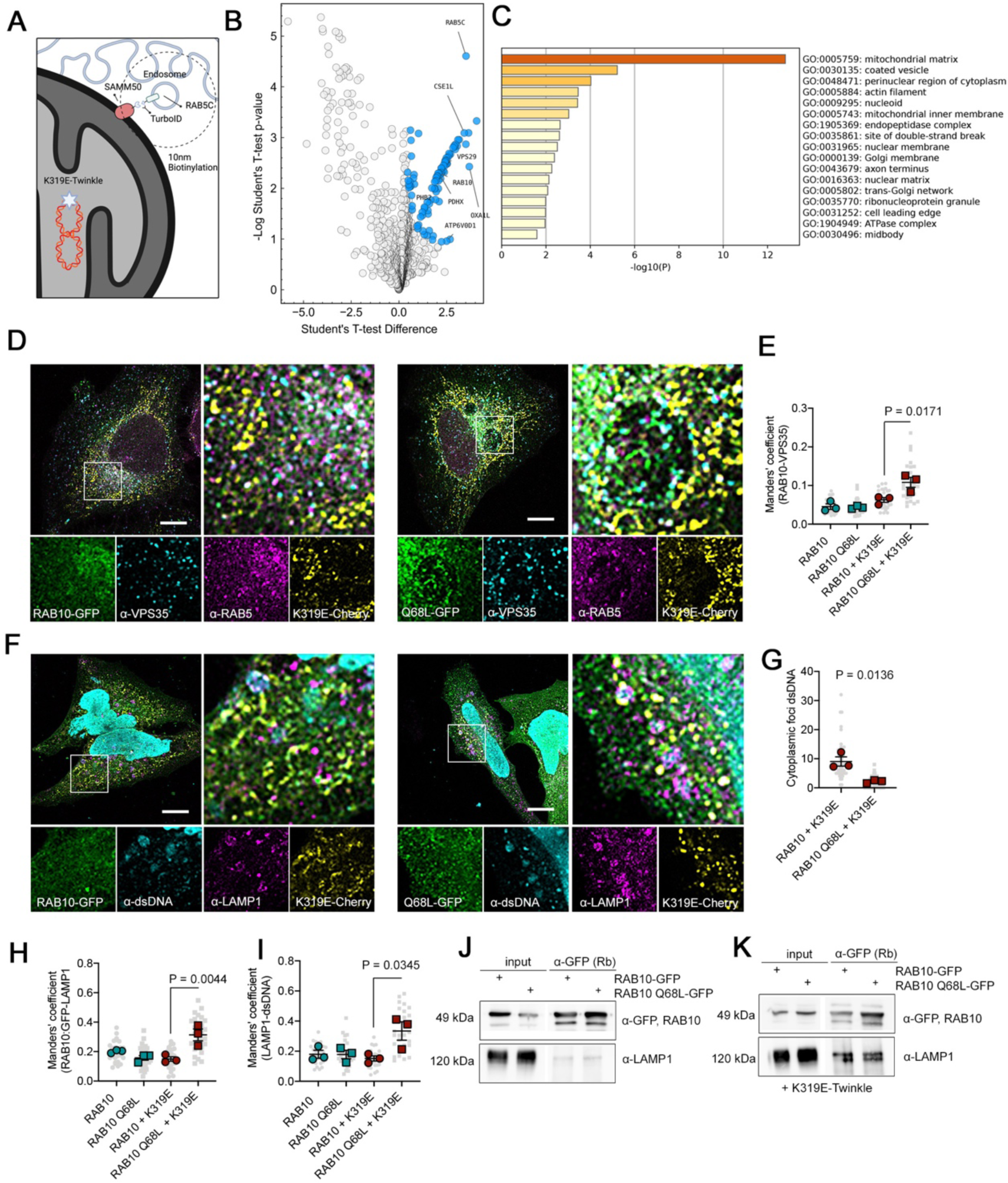
mtDNA damage follows a degradation pathway related to lysosomes. (**A**) Graphical model for the proximity proteomics approach. TurboID was split and combined with the mitochondrial outer membrane protein SAMM50 and the early endosomal marker RAB5C (biorender.com) (**B**) Volcano Plot showing proteins enriched after biotinylation and purification of cells expressing SplitTurboID plasmids and K319E-Cherry. Differentially expressed proteins compared with cells transduced with SAMM50-SplitCt (significant: q-value < -0.05 and absolute log2 fold change>1) are highlighted in blue (n=6). (**C**) Pathway enrichment analysis with Metascape showing GO terms for proteins differentially enriched. (**D**) Immunostaining and (**E**) quantification of HeLa cells expressing RAB10-GFP, RAB10^Q68L^-GFP and TWNK^K319E^-mCherry plasmids and labeled with α-VPS35 and α-RAB5, and, (**F**) cells labeled with α-dsDNA and the lysosomal marker α-LAMP1 (n=3, 10 images per replicate). (**G**) Quantification of cytosolic dsDNA foci in cells expressing RAB10-GFP, RAB10^Q68L^-GFP and TWNK^K319E^-Cherry plasmids (n=3, <15 cells per replicate). (**H**) Manderścorrelation coefficient between RAB10-GFP and LAMP1, and (**I**) LAMP1 and dsDNA (n=3, 10 images per replicate). **j**) RAB10-GFP co-Immunoprecipitation in steady state and (**K**) in cells expressing TWNK^K319E^-Cherry with the lysosomal protein LAMP1. *P* values were calculated using One-way ANOVA with Tukey correction for multiple comparison. Scale bar, 10 μm. Data is presented as mean ± SEM

To isolate high levels of biotinylated proteins, we generated HEK293 stable cell lines by lentiviral transduction. Western blot analysis of biotinylation revealed residual activity in cells expressing only SAMM50-SplitTurboCt-HA (Fig. S2C). Then, to minimize potential background interference, we used these cells as a negative control, and cells expressing both SAMM50-SplitTurboCt-HA and RAB5C-SplitTurboNt-V5 as the experimental approach. To specifically interfere with mtDNA replication, we expressed a dominant negative mutation of the mitochondrial helicase Twinkle (TWNK^K319E^). The equivalent mutation in mice (TWNK^K320E^) has been validated as a reliable tool to induce mtDNA alterations, oxidative stress, and replication defects^11–13^. Expression of this missense mutation in HeLa cells resulted in mild mitochondrial fragmentation but did not interfere with cell viability (Fig. S1D and E).

The proximity proteome of the mitochondria and endosomes at steady state showed enrichment of proteins involved in intra-Golgi vesicle transport and mitochondrial protein import (Fig. S1F and G; Supplementary Table S1). On the contrary, in cells with mtDNA replication stress, we observed a complete rewiring of the mitochondrial-endosome proteome (Figure 1B). Pathway analysis revealed the presence of mitochondrial matrix proteins (galactokinase, GALK1; pyruvate dehydrogenase component, PDHX; fumarate hydratase, FH), nucleoid and mitochondrial inner membrane proteins (DNAJC11; prohibitin 2, PHB2; OXA1L), pointing to a rupture of the mitochondrial outer membrane and hence, biotinylation of mitochondrial matrix components (Figure 1C). In addition, we also detected proteins related to coated vesicles such as the retromer subunit VPS29 and RAB10, together with the lysosomal proteins MYO1B, MYO6, ATP6V0D1 and cathepsin D (CTSD) (Figure 1C; Supplementary Table S1). Thus, mtDNA damage changes the proximity proteome of mitochondria towards lysosomal recruitment and degradation of mitochondrial components.

### RAB10 is recruited to late endosomes for the elimination of leaked cytosolic mtDNA

The small GTPase RAB10 participates in mitochondrial quality control. Upon mitochondria depolarization, RAB10 binds the autophagy receptor optineurin and promotes its accumulation, facilitating mitophagy^14^. We sought to investigate the subcellular localization of RAB10 upon mtDNA replication stress. The expression of RAB10-GFP or the gain-of-function mutation, RAB10^Q68L^-GFP, did not affect mitochondrial morphology significantly (Fig. S2A and C). However, cells co-expressing TWNK^K319E^ suffered from an exacerbated mitochondrial fragmentation (Fig. S2B and D). RAB10-GFP and RAB10^Q68L^-GFP presented a reticular pattern, colocalizing with the endoplasmic reticulum (CNX, Calnexin; Fig. S3A and B). In these cells, the association of RAB10-GFP with the retromer was enhanced (Figure 1D and E). Noteworthy, we noted the presence of accumulated cytoplasmic DNA foci, not resembling the typical mtDNA staining (Figure 1F; arrows). Although these foci were present in the majority of TWNK^K319E^-cells, their number was lower in cells co-expressing the gain-of-function allele of RAB10 (Figure 1G). Further immunostaining showed that late endosomal marker LAMP1 decorated these cytosolic DNA foci together with RAB10 (Figure 1F). Colocalization between RAB10 and LAMP1, as well as between LAMP1 and dsDNA, was remarkably enhanced in RAB10^Q68L^-GFP cells (Figure 1H and I). Pull-down experiments with RAB10-GFP confirmed a strong association with LAMP1 upon mtDNA replication stress (Figure 1J and K). On the contrary, we observed very weak co-IP of RAB5, neither with RAB10, nor with LAMP1 or VPS35, both in control cells and TWNK^K319E^-cells (Fig. S3D). Altogether, these data indicate that mtDNA instability triggers the reorganization of the vesicular system. mtDNA leaked induced by mtDNA replication stress induces translocation of RAB10 from the ER to late endosomes, which are transported to cytosolic regions containing DNA for degradation. RAB10 participates in lysosomal recruitment, mitochondrial fragmentation and removal of leaked mtDNA.

### The retromer facilitates the release of mitochondrial matrix components during mitochondrial stress

One of the mechanisms proposed for mtDNA elimination is the formation of MDVs. The retromer, a heterotrimer formed by VPS26, VPS35, and VPS29, is the main protein complex in charge of retrograde vesicle transport through the endosome-lysosomal pathway^15^. The core component VPS35 contributes to MDVs formation directed to peroxisomes^16^. Our proteomic approach detected, upon mtDNA replication stress, enrichment on the retromer subunit VPS29. Nevertheless, no genetic variation in *VPS29* influencing the function of the retromer complex has been yet identified. On the contrary, mutations in the retromer core component *VPS35* interfere with the functionality of the complex. In particular, the Parkinsońs disease-associated mutation p.D620N restricts cargo selection and mitochondrial clearance^17, 18^. Thus, we analyzed the formation of MDVs upon mtDNA replication stress and the expression of VPS35 and VPS35^D620N^-GFP.

Among the different types of MDVs, vesicles carrying pyruvate dehydrogenase (PDH), and lacking the outer membrane protein TOM20, are bona-fide markers for vesicles containing mitochondrial matrix cargo^19^. At steady state, expression of both versions of VPS35-GFP induced a slight increase of filamentous mitochondria (Fig. S2E and G). VPS35 did not colocalize with the mitochondrial marker PDH but with the early endosomal marker RAB5 (Fig. S3E and F). Colocalization between VPS35-GFP and dsDNA was also not observed (Fig. S3G).

Expression of TWNK^K319E^ in combination with VPS35 showed increased fragmentation of the mitochondrial network (Fig. S2F and H). In this context, we detected the formation of PDH^+^/TOM20^-^ particles (Figure 2A and B). However, these particles were only detected when lysosomal activity was blocked with chloroquine. We also found VPS35 in the exit sites of PDH (Figure 2C and D). Remarkably, PDH^+^/TOM20^-^ particles were not present in cells expressing VPS35^D620N^-GFP (Fig 2A and B).

**Figure 2.**
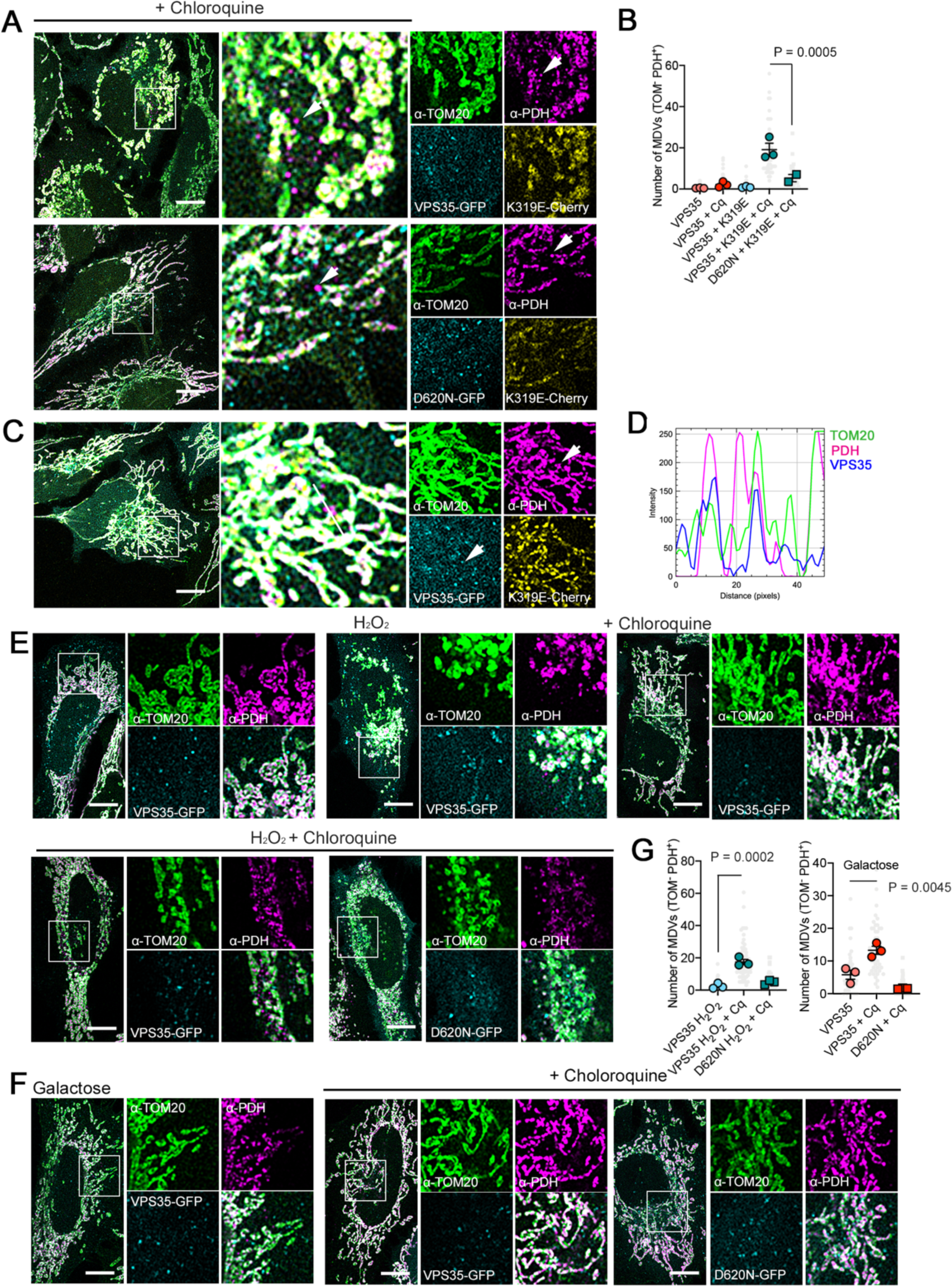
mtDNA stress and oxidation initiate release of mitochondrial components. (**A**) HeLa cells expressing VPS35-GFP, TWNK^K319E^-Cherry and labelled with the mitochondrial outer membrane marker α-TOM20 and mitochondrial matrix α-PDH. (**B**) Quantification of Mitochondrial derived vesicles (MDVs; α-TOM20 positive, α-PDH negative; n=3, >15 cells per replicate). (**C**) Immunostaining and (**D**) fluorescence profile of cells expressing VPS35-GFP and labelled with α-TOM20 and α-PDH. Fluorescence intensity profile was obtained from 50-pixel line labelled in C. (**E**) MDVs immunostaining in cells were treated with 200 μM H_2_O_2_ 4h prior to fixation or (**F**) grown in Galactose medium overnight. (**G**) Quantification of Mitochondrial derived vesicles in H_2_O_2_ and Galactose treated cells. Where indicated, cells were treated with 10 μM Chloroquine 4h prior fixation to block lysosomal function. (n=3, >15 cells per replicate). *P* values were calculated using One-way ANOVA with Tukey correction for multiple comparison. Scale bar, 10 μm. Data is presented as mean ± SEM

Next, we asked if the elimination of mitochondrial content mediated by VPS35 was also occurring when interfering with mitochondrial function by other means. Given that mtDNA replication stress causes oxidative damage^6^ and this is one of the most common forms for natural acquisition of mtDNA mutations^20^, we treated our cells with a sublethal concentration of H_2_O_2_. Again, we observed the release of mitochondrial matrix content only when the lysosomal function was blocked with chloroquine (Figure 2E and G). In addition, MDVs were also observed when cells were grown in galactose instead of glucose, a condition that supports mitochondrial metabolism and oxidation^21^ (Figure 3F and G). This mechanism was non-functional for the PD-associated mutation VPS35^D620N^. These data confirm that upon mtDNA replication stress and in an oxidative environment, the mitochondrial matrix material is eliminated through MDVs supported by VPS35.

**Figure 3.**
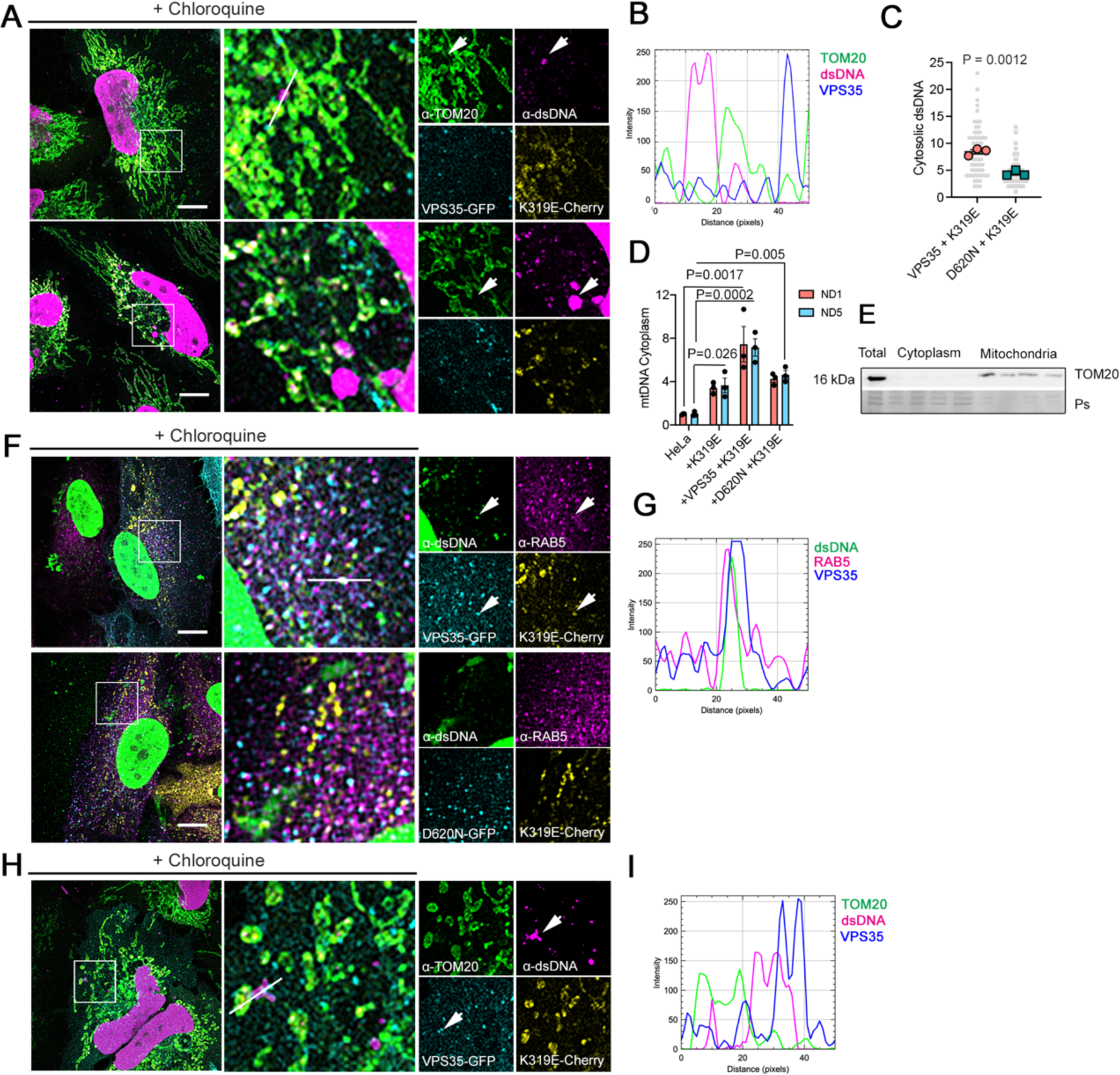
VPS35 enhances extraction of mtDNA. (**A**) HeLa cells expressing TWNK^K319E^-Cherry and VPS35-GFP or VPS35^D620N^-GFP, labelled α-TOM20 and α-dsDNA. (**B**) Fluorescence profile of 50-pixel line labelled in A. (**C**) Quantification of cytoplasmic dsDNA foci (dsDNA^+^-TOM20^-^). (n=3, >15 cells per replicate) (**D**) qPCR quantification of *ND1* and *ND5* mtDNA genes in mitochondria-free cytosolic fractions. (n=3) (**E**) Western blot analysis in cells treated with digitonin to eliminate mitochondria and used for qPCR. Ps, Ponceau. (**F**) Immunostaining and (**G**) 50-pixel line fluorescence profile of cells expressing VPS35-GFP and labelled with α-RAB5 and α-dsDNA. (**H**) α-TOM20 and α-dsDNA immunostaining and (**I**) fluorescence profile of mtDNA exit site. Where indicated, cells were treated with 10 μM Chloroquine 4h prior fixation to block lysosomal function. *P* values were calculated using StudentśT-test (C) or One-way ANOVA with Tukey correction for multiple comparison (D). Scale bar, 10 μm. Data is presented as mean ± SEM

### mtDNA ejection during mtDNA stress occurs directly in VPS35 late endosomes

We sought to investigate if mtDNA extraction upon mtDNA stress follows the same path as observed for mitochondrial matrix components. The presence of cytosolic dsDNA foci was observed by confocal microscopy in cells expressing TWNK^K319E^ and VPS35 (Figure 3A and B). VPS35^D620N^ cells, however, showed a reduced number of extramitochondrial dsDNA (Figure 3C). To confirm the presence of mtDNA in the cytoplasm, we performed a qPCR analysis of mtDNA genes in mitochondria-free cytosolic fractions. In agreement, we found an increase of both *MT-ND1* and *MT-ND5* in cells with mtDNA replication stress (Figure 3D and E). Expression of VPS35 further increased extraction, while no changes were observed in cells with the dominant negative VPS35^D620N^. In this context, VPS35 could be also detected together with RAB5 particles containing dsDNA (Figure 3F and G). Similar to the mitochondrial matrix MDVś, we found VPS35-GFP in the exit sites of the mtDNA (Figure 3H and I).

Then, we performed correlative light-electron microscopy (CLEM) to visualize the specific location of cytosolic mtDNA foci and combined it with electron tomography to gain volume information. We stained chloroquine-treated cells with SYBR Gold for DNA and PK Mito Deep Red for mitochondria. Using this approach, no DNA was found outside the mitochondrial compartment in control cells (Fig. S4A). In contrast, cells expressing TWNK^K319E^ showed cytosolic DNA foci (Figure 4A; Fig. S4B). These cells also had reduced mitochondrial size and fewer cristae per mitochondria (Fig. S4C and D). The correlation of confocal images with transmission electron microscopy and electron tomogram showed that areas containing mtDNA were enriched in recycle organelles at different stages of maturation. Cytosolic DNA was encapsulated in a recycling organelle (Figure 4B and C; Fig. S4F). Noteworthy, none of all cells analyzed with CLEM (n=6) showed DNA inside a small vesicle. Then, we performed the same approach in cells expressing TWNK^K319E^ and VPS35-CFP (Figure 4D; Fig. S4G). Correlation experiments confirmed the presence of VPS35 in the recycling organelles in close apposition with mitochondria (Figure 4E and G). In agreement with our confocal data (Figure 3C), electron tomography reconstitution of the mtDNA exit site confirmed the transfer of mtDNA to a nearby vesicle labeled also with VPS35-CFP (Figure 4F and H). These data suggest that for the mitochondrial genome, the elimination occurs directly to recycling organelles labeled with VPS35. Degradation of cytosolic mtDNA is impaired in cells expressing the Parkinsońs disease-associated mutation VPS35^D620N^.

**Figure 4.**
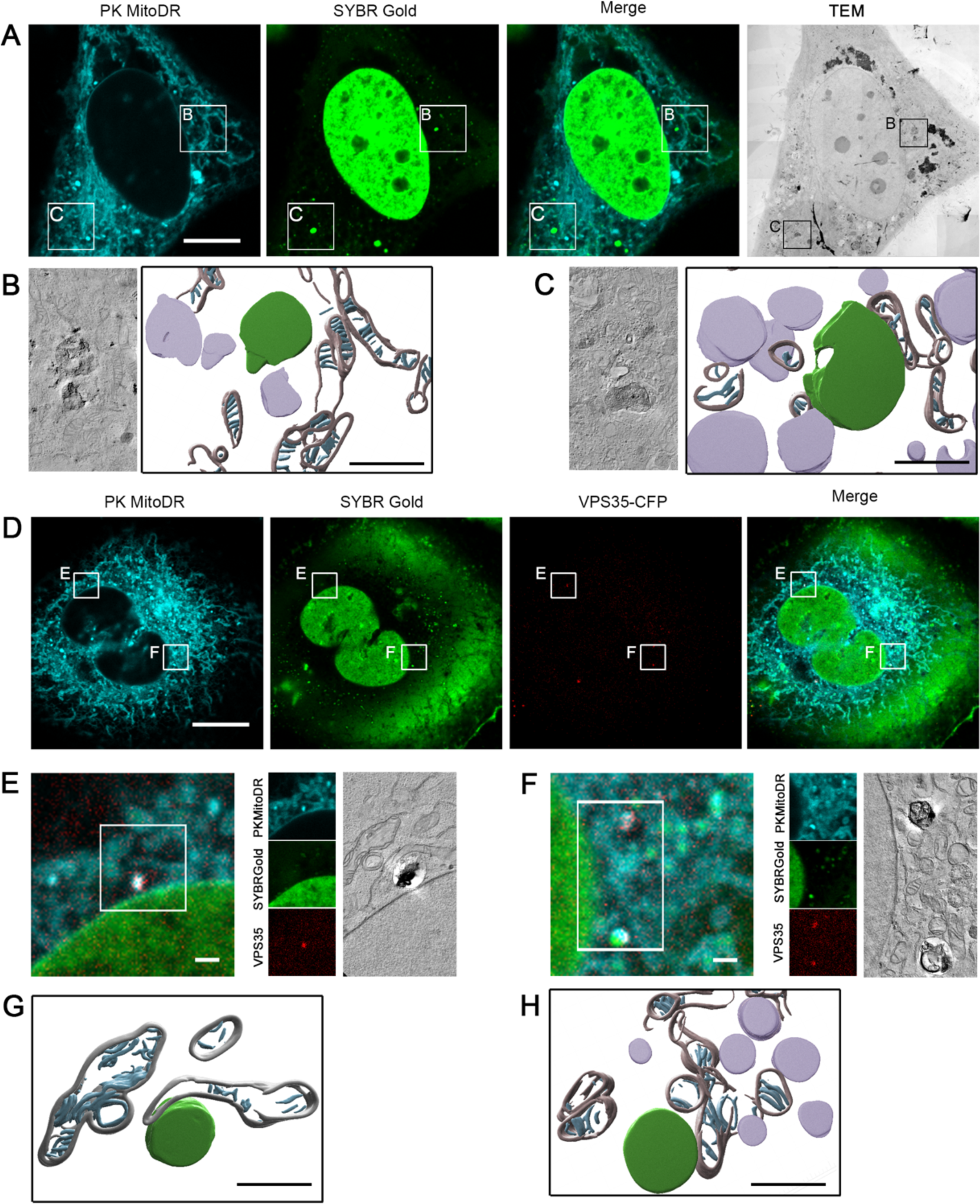
Cytoplasmic mtDNA localize with VPS35 recycling organelles. (**A**) Correlative light-electron microscopy (CLEM) of cells expressing TWNK^K319E^-Cherry and labelled with SYBR Gold and PK MitoDeep Red. (**B, C**) Volumetric reconstitution of electron tomographies of cytosolic mtDNA areas. Lysosome containing DNA is shown in green, recycling organelles in lilac, mitochondrial outer membrane in grey and mitochondrial cristae in cyan. (**D**) TWNK^K319E^-Cherry cell further expressing VPS35-CFP indicating the areas used for CLEM. (**E**) Correlation of VPS35-CFP foci and (**F**) mtDNA exit site. (**G, H**) Volumetric reconstitution of electron tomographies in VPS35-CFP cells. In all cases, cells were treated with 10 μM Chloroquine and loaded for 1h with the indicated dyes prior fixation. Scale bar 500 nm for reconstituted tomograms (B, C, G and H), 1 μM (E, F) and 10 μM (A, D).

### Generation of long mtDNA deletion in *Drosophila* larval epidermis

Our findings suggest that the retromer enhances mitochondrial quality control in response to mtDNA damage. We hypothesized that overexpression of VPS35 might be beneficial *in vivo*, by reducing mitochondrial burden associated with mtDNA mutations. To investigate our hypothesis, we used a well-established GAL4/UAS *Drosophila* model based on nucleases^22^. *Drosophila* mtDNA harbors two target sites for the restriction enzyme AflIII. Co-expression of mitochondria-targeted AflIII (mitoAflIII) and the DNA ligase from phage T4 (mitoT4lig) (Figure 5A; *UAS-mitoAflIII, UAS-mitoLigase*), results in the generation of 2564 b.p deletion in the mtDNA, including *CytB* gene (mtDNA^Δ10.789-13.372^).

**Figure 5.**
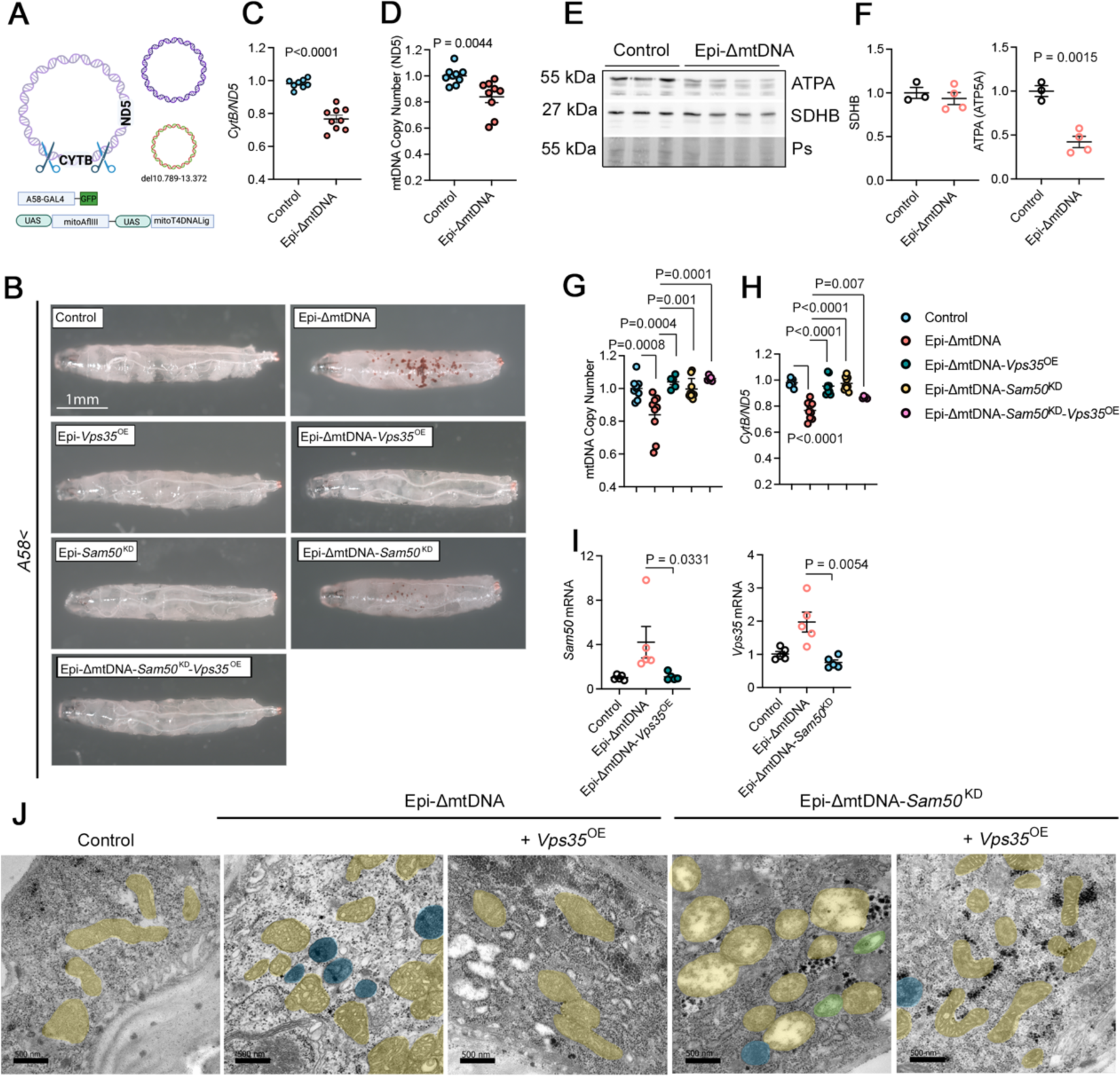
VPS35 overexpression recovers mitochondrial defects associated to mtDNA deletions in *Drosophila*. (**A**) Schematic representation of the approach to generate mtDNA deletion covering 2.564 b.p in larval epidermis. The restriction enzyme AflIII and the T4-DNA Ligase directed to mitochondria were both expressed under UAS promotor. A-58 GAL4 promotor was used to drive the expression of the transgenes in the larval epidermis. (**B**) *CytB/ND5* ratio showing mtDNA heteroplasmy obtained by qPCR from total DNA extracts of L3 larvae. (Control, n=8; Epi-ΔmtDNA, n=9). (**C**) mtDNA copy number quantification using *ND5* gene and *TUBB* as a mitochondrial and nuclear gene respectively (n=9). (**D**) Western blot analysis and (**E**) quantification in total protein extracts of L3 larvae for SDHB ATPA. Ponceau S (Ps) was used as a loading control. (Control, n=3; Epi-ΔmtDNA, n=4). (**F**) Bright field images of transgenic L3 larvae used in this study. (**G**) mtDNA copy number and (**H**) mtDNA heteroplasmy quantification upon genetic manipulations. (Control, n=9; Epi-ΔmtDNA, n=9; Epi-ΔmtDNA-*Vps35*^OE^, n=5; ΔmtDNA-*Sam50*^KD^, n=9; ΔmtDNA-*Vps35*^OE^-*Sam50*^KD^, n=5). (**I**) Relative levels of *Sam50* and *Vps35* mRNA in Epi-ΔmtDNA larvae. (n=5 per genotype). (**J**) Electron microscopy images of larval epidermis. Mitochondria was pseudo-colored in yellow, lysosomes in blue (dark content) and autophagosomes (recycling organelle with double membrane) in green. *P* values were calculated using StudentśT-test (B, C, E) and One-way ANOVA with Tukey correction for multiple comparison (G, H, I). Scale bar, 1 mm (F) and 500nm (J). Data is presented as mean ± SEM.

Expression of both UAS contracts (recombined *UAS-mitoAflIII, UAS-mitoLigase*) by the majority of tested GAL4 drivers (neurons, epithelial, muscles, or fat bodies), leads to embryonic or early larval lethality^22^. Consistently, upon recombination mediated by the larval epidermal driver A58 at 25 °C (*A58-GAL4 >UAS-mitoAflIII,UAS-mitoLigase;* from now on Epi-ΔmtDNA), only a few larvae with massive brown melanized foci could survive till pupal stage. To overcome the lethality, we maintained the crosses at 18°C. At this temperature, the GAL4 activity is at its lowest levels, leading to larval survival till the pupal stage^23^. At this temperature, the chromogenic foci were more restricted and larvae could pupate, although only ∼1% of pupae emerged (Figure 5B). Hence, this temperature and L3 larvae were selected for further experiments.

Expression of mitoAflIII was confirmed to peak in the early L3 instar stage and decay in the mid-stage (L3-E Vs. L3-M; Fig. S5A). Conventional PCR using oligos flanking mtDNA^Δ10.789-13.372^ confirmed the presence of the mtDNA deletion in Epi-ΔmtDNA (Fig. S5B). Accordingly, the *CytB*/*ND5* ratio was reduced (Figure 5C) and the mtDNA copy number decreased to a bigger extent when *CytB* was used for mtDNA copy number quantification (Figure 5D; Fig. S6A). Western blot analysis from total protein extracts showed equal levels of the mitochondrial complex II subunit succinate dehydrogenase B (SDHB) and depletion of complex V subunit ATPA (Figure 5E and F). Given that mitochondrial complex II is encoded by nuclear genes and complex V by both nuclear and mitochondrial genomes, this result confirmed that mtDNA depletion was not related to reduced mitochondrial mass but indeed due to a lower number of mtDNA copies. Thus, the co-expression of mitoAflIII and mitoT4lig in the epidermis of *Drosophila* larvae efficiently deletes mtDNA and increases mtDNA heteroplasmy.

### VPS35 promotes mtDNA clearance *in vivo*

To test if *Vps35* elevation supports mtDNA clearance *in vivo*, we overexpressed *Vps35* in the larval epidermis by crossing the UAS-Vps35 construct with the A58-GAL4 driver (Epi-*Vps35*^OE^). Epi-*Vps35*^OE^ larvae did not exhibit macroscopic abnormalities and there were no observable morphological or developmental changes (Figure 5B). qPCR quantification of total RNA extracts revealed a 9-fold increase in *Vps35* mRNA (Fig. S6B). Additionally, Epi-*Vps35*^OE^ larvae had a moderate but significant increase in mtDNA copy number (Fig. S6C). Next, we generated Epi-ΔmtDNA*-Vps35*^OE^. In this background, *Vps35* mRNA levels were increased 5-fold on average compared to *Epi-ΔmtDNA* (Fig. S6D). We did not observe major macroscopic and morphological abnormalities in larvae and the chromogenic foci initially observed in Epi-ΔmtDNA larvae disappeared (Figure 5B). Quantification of mtDNA copy number and *CytB/ND5* ratio showed normalized mtDNA copies in Epi-ΔmtDNA-*Vps35*^OE^ compared to wt larvae (Figure 5G and H). Moreover, western blot data confirmed higher levels of the complex V protein ATPA and no changes in the nuclear-encoded complex II SDHB (Fig. S5E and F). All this data indicates that *in vivo*, a high level of *Vps35* is sufficient to reduce damage in the mitochondrial genome and restore normal mtDNA levels.

### VPS35 genetically interacts with SAM50 *in vivo*

We previously described that localization of mtDNA in VPS35 endosomes upon mtDNA damage in cells is dependent on the mitochondrial outer membrane protein SAMM50^6^. To validate this observation *in vivo*, we generated SAMM50 knockdown in the epidermis of larvae. In *Drosophila*, SAMM50 has not been characterized, although the gene *CG7639* (from now on, *Sam50*) was predicted to be the human ortholog. Therefore, we knockdown *Sam50* in the epidermis of larvae by crossing *A58-GAL4* with the *UAS-Sam50*-RNAi construct (Epi-*Sam50*^KD^). qPCR mRNA quantification showed up to 60% reduction in *Sam50* mRNA (Fig. S6G) and a slight reduction of mtDNA copy number (Fig. S6H), with no obvious abnormalities (Figure 5B). In contrast to what we observed before, western blot analysis showed reduced nuclear encoded SDHB, suggesting that, in this case, the mtDNA copy number decrease was related to reduced mitochondrial mass (Fig. S6I and J).

To investigate the Sam50 function in mtDNA damage epidermis and for better Sam50 knockdown efficiency, we generate a homozygous *Sam50*^KD^ in Epi-ΔmtDNA background (Epi-ΔmtDNA-*Sam50*^KD^). In this case, melanized foci were reduced compared to Epi-ΔmtDNA but still present (Figure 5B). *Sam50* mRNA levels decayed up to 80% (Fig. S6K). Surprisingly, a significant reduction of *Sam50* also restored mtDNA copy number and *CytB*/*ND5* ratio in Epi-ΔmtDNA larvae (Figure 5G and H).

mRNA quantification by qPCR showed that both *Vps35* and *Sam50* were upregulated in Epi-ΔmtDNA larvae (Figure 5I). *Vps35* upregulation was restored upon *Sam50* KD in ΔmtDNA genetic background and vice versa (Figure 5I). To investigate *Vps35* and *Sam50* interaction upon mtDNA damage, we performed an epistatic experiment, upregulating *Vsp35* while also reducing *Sam50* in Epi-ΔmtDNA (Epi-ΔmtDNA-*Sam50*^KD^-*Vps35*^OE^) (Fig. S6L). In these larvae, melanized foci were cleared (Figure 5B) and mtDNA copy number was still recovered (Figure 5F and G) but *CytB/ND5* ratio slightly changed compared to Epi-ΔmtDNA (Figure 5H).

Finally, we investigated mitochondria ultrastructure changes triggered by these genetic modulations. Electron microscopy preparation of the larvae epidermis showed cristae aberrations in the epidermis of Epi-ΔmtDNA (Figure 5J). In addition, we noticed the accumulation of lysosomes and multivesicular bodies intercalated with mitochondria (Figure 5J, blue pseudo-colored). Epi-*Vps35*^OE^ in Epi-ΔmtDNA background showed complete recovery of mitochondrial ultrastructure similar to control larvae. In contrast, mitochondria from Epi-ΔmtDNA-*Sam50*^KD^ were swollen and completely lacking cristae (Figure 5J). Epidermis from Epi-ΔmtDNA-*Sam50*^KD^ was also characterized by the accumulation of autophagosomes (green pseudo-colored). However, Epi-ΔmtDNA-*Sam50*^KD^-*Vps35*^OE^ showed similar mitochondrial morphology to Epi-ΔmtDNA but a lack of autophagy organelles, albeit the accumulation of pycnotic granules in the cytosol (Figure 5J).

Altogether, these data indicate that both *Sam50* and *Vps35* participate in the genetic response activated upon mtDNA damage and interestingly with an epistatic interaction between them in response to mtDNA damage.

### VPS35 overexpression inhibits the activation of bulk mitophagy

*Vps35* overexpression has a beneficial effect on the changes induced by mitochondrial impairment. However, these effects are also influenced by *Sam50* mRNA level. To clarify the role of these two proteins we shifted to cells and used Hela doxycycline-inducible SAMM50 KD cells^24^. In these cells, *SAMM50* KD induces mtDNA leakage and robust mitophagy response^25, 26^. Hence, we used this background for VPS35 overexpression.

5 days of constant doxycycline supply was enough to strongly reduce SAMM50 protein level (Figure 6A and B). However, the autophagy receptor SQSTM1/p62, the mitophagy receptor optineurin, and the lipidated form of the autophagosome protein LC3B (LC3B-II) were unchanged (Figure 6A and B). mtDNA copy number was also unchanged after 5 days (Figure 6C). Super-resolution microscopy with STED and Airyscan2 showed a rupture of the mitochondrial inner membrane and mtDNA release into the cytoplasm (Figure 6D and E). Consequently, we detected an early activation of the innate immune response (Figure 6F). Prolonged doxycycline supply for 10 days, elicited a strong mitophagy response, illustrated by upregulation of LC3B-II and optineurin, depletion of SQSTM1/p62, and mtDNA copy number depletion (Figure 6A, B, and C).

**Figure 6.**
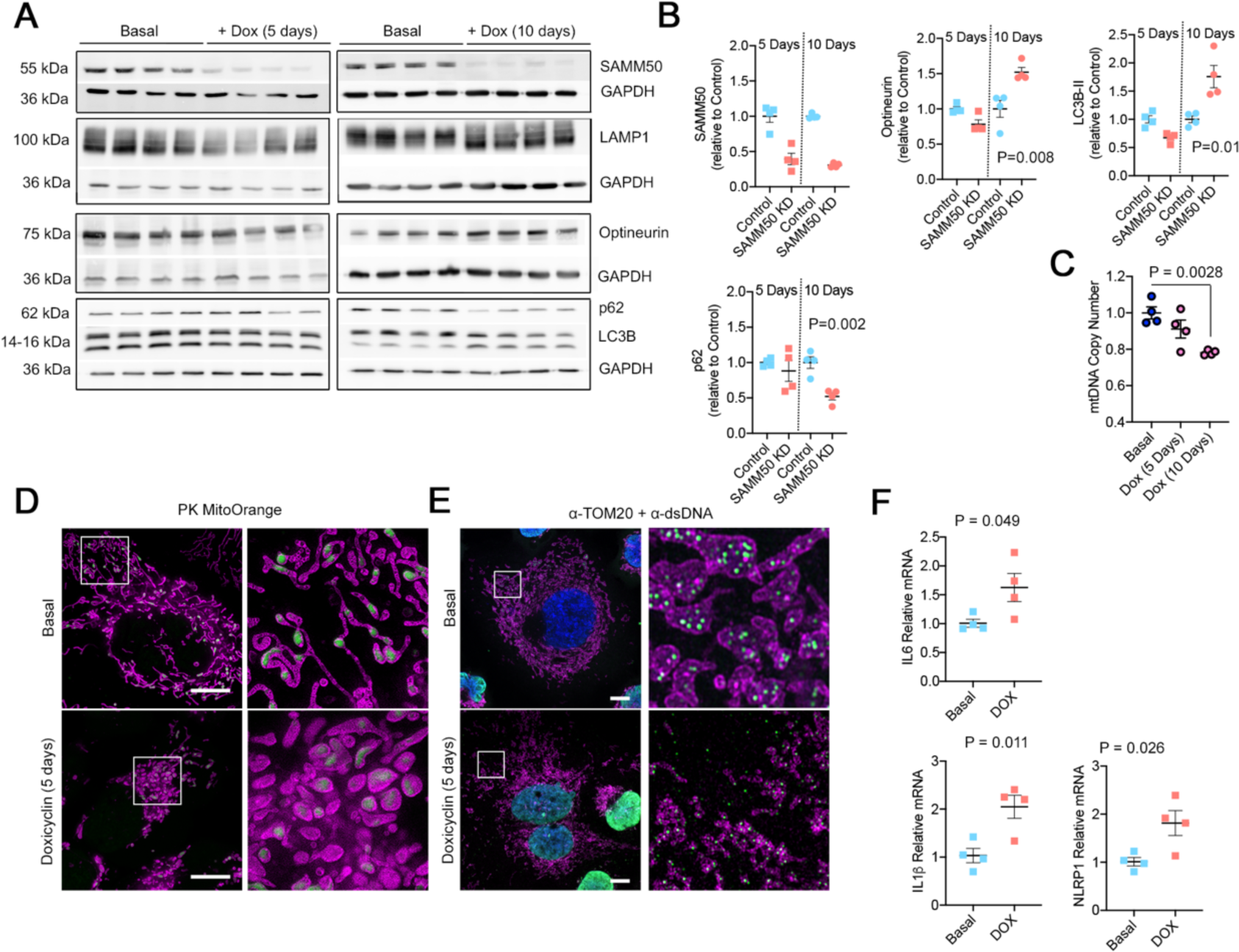
SAMM50 KD triggers mtDNA loss, activation of macroautophagy and the innate immune response. (**A**) Western blot analysis and (**B**) quantification in control and *SAMM50* KD for autophagy/mitophagy markers at the indicated time points after KD induction. α-GAPDH was used as a loading control. (n=4). (**C**) mtDNA copy number quantification after *SAMM50* KD (n=4). (**D**) Super resolution microscopy in cells loaded with the mitochondrial inner membrane marker PK MitoOrange. (**E**) Airy Scan super resolution microscopy of *SAMM50* KD cells labelled with α-TOM20 and α-dsDNA. (**F**) mRNA quantification of innate immune related genes. *GAPDH* mRNA was used for normalization (n=4). *P* values were calculated using StudentśT-test (B, F) and One-way ANOVA with Tukey correction for multiple comparison (C). Scale bar, 10 μm. Data is presented as mean ± SEM.

Then, we transduced inducible SAMM50 KD HeLa cells with lentivirus containing *VPS35-V5* cDNA. Already in a basal state, these cells showed an increase in the lipidated form of LC3B and SQSTM1/p62, with no changes in other components of the vesicular system (Figure 7A and B). Block of lysosomal activity by chloroquine revealed an accumulation of the lysosomal protein LAMP1 in cells overexpressing VPS35 (Figure 7C and D), suggesting increased lysosomal turnover. In this case, chronic downregulation of *SAMM50* for 10 days did not change the autophagy/mitophagy profile (Figure 7E and F). VPS35 overexpression did not modify the mtDNA copy number neither in the steady state nor after *SAMM50* KD (Figure 7G and H). Immunostaining in cells overexpressing VPS35 showed an increase in lysosome-mitochondria colocalization upon SAMM50 KD and in cells overexpressing VPS35 already in a steady state (Figure 7I and J). However, this was not sufficient to eliminate cytosolic mtDNA induced by SAMM50 KD and counteract the activation of the innate immune response (Figure 7K and L), confirming that SAMM50 is essential to locate mtDNA inside lysosomes. Nevertheless, *VPS35* overexpression enhances autophagy flux by increasing lysosomal content and turnover. Hence, VPS35 overexpression accelerates the turnover of damaged mitochondria upon SAMM50 KD preventing the activation of a robust mitophagy response.

**Figure 7.**
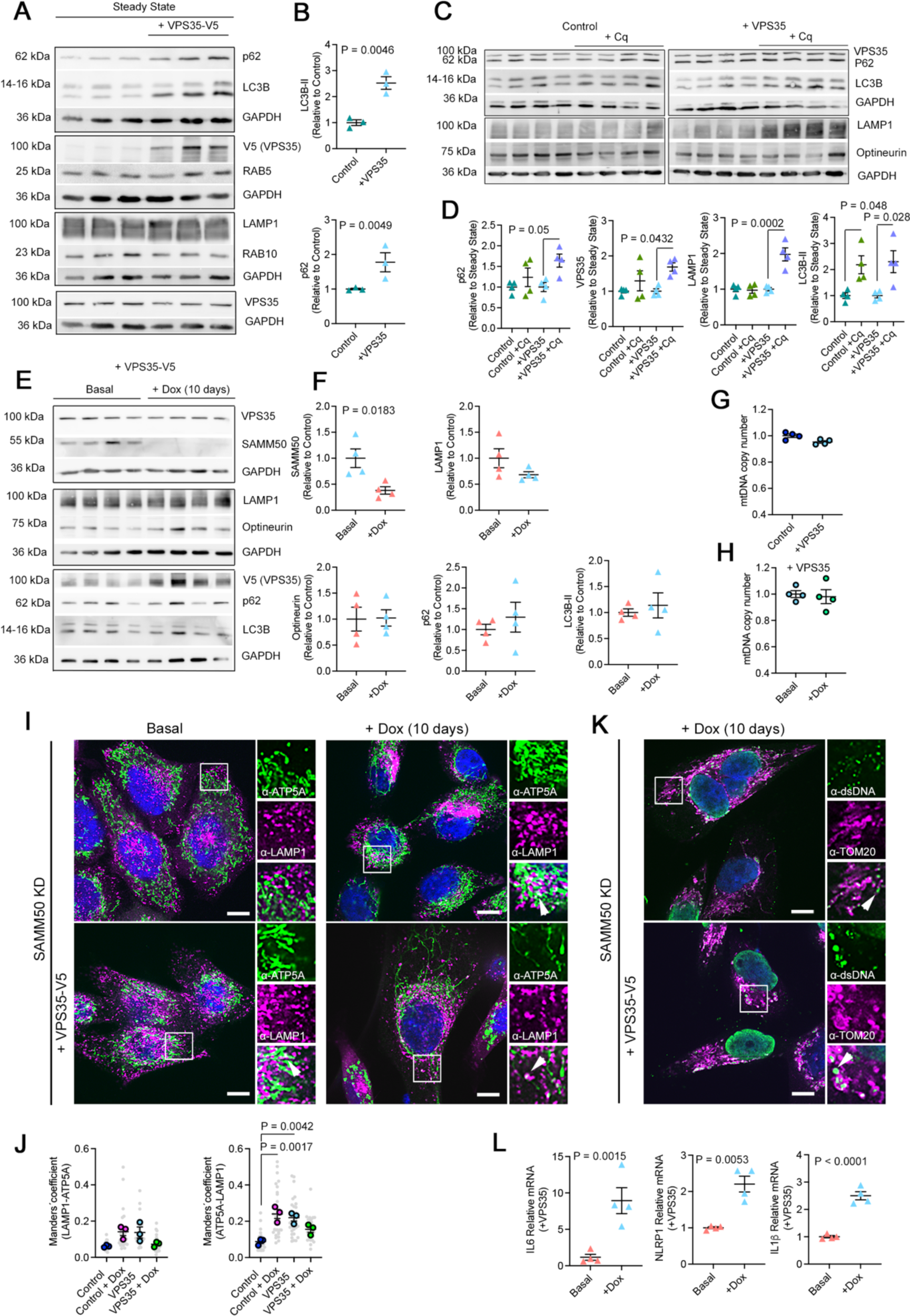
VPS35 overexpression accelerates autophagy flux through lysosomal accumulation. (**A**) Western blot analysis and (**B**) quantification of macroautophagy and proteins from the vesicular system in control and cell expressing VPS35-V5 (n=3). (**C**) Western blot and (**D**) quantification in cells treated with 10 μM Chloroquine (Cq) for 4h to assay autophagy flux (n=4). (**E**) Western blot and (**F**) quantification in SAMM50 knock-down cells overexpressing VPS35-V5. α-GAPDH was used as a loading control (n=4). (**G**) mtDNA copy number quantification for control and VPS35 cells in steady state and (**H**) upon SAMM50 KD (+ Dox) (n=4). (**I**) α-ATP5A and α-LAMP1 immunostaining in basal and upon SAMM50 KD (+ Dox) in VPS35-V5 cells. Arrows depict colocalization foci. (**J**) Manderś correlation coefficient between LAMP1 and ATP5A and vice versa (n=3, 10 images per replicate) (**K**) α-dsDNA and α-TOM20 immunostaining in VPS35-V5 and SAMM50 KD cells. (**L**) mRNA quantification of innate immune related genes. *GAPDH* mRNA was used for normalization (n=4). *P* values were calculated using Studentś T-test (B, L) and One-way ANOVA with Tukey correction for multiple comparison (J). Scale bar, 10 μm. Data is presented as mean ± SEM.

### Elevated VPS35 reverses transcriptome profile in ΔmtDNA larval epidermis

For an unbiased and genome-wide insight overview of different pathways activated upon ΔmtDNA, we conducted a transcriptome analysis in isolated epidermis from L3 ΔmtDNA larvae (Fig. S7A and B). As expected by the presence of the deletion, mtDNA^Δ10.789-13.372^, mitochondrial mRNAs for *CytB*, *ND1* and *lrRNA*, genes included in the deletion, were strongly reduced (Fig. S7C). Then, we performed differential expression analysis using four independent bioinformatic methods (EdgeR-QFL, EdgeR-LRT, limma-voom and DESeq2) and identified 541 genes with significant upregulation, while 311 genes were downregulated (Figure 8A; Supplementary Table S2). Pathway analysis with DAVID and Gene Set Enrichment Analysis (GSEA) with Gene Ontology pathways uncovered upregulation of proteolysis, innate immune response, and redox control routes (Figure 8B and C; Fig. S7D and E. Supplementary Table S3). To further validate these data, we selected some upregulated targets and performed qPCR analysis in independent RNA isolates. We found consistent enrichment of the Glutathione GST transferase genes *GstD2*, *GstD5*, *GstD9*, *GstE5*, *GstE7* and *Gst9,* along with the mitochondrial proteases *Spg7* and *Afg3l2* (Figure 8D).

**Figure 8.**
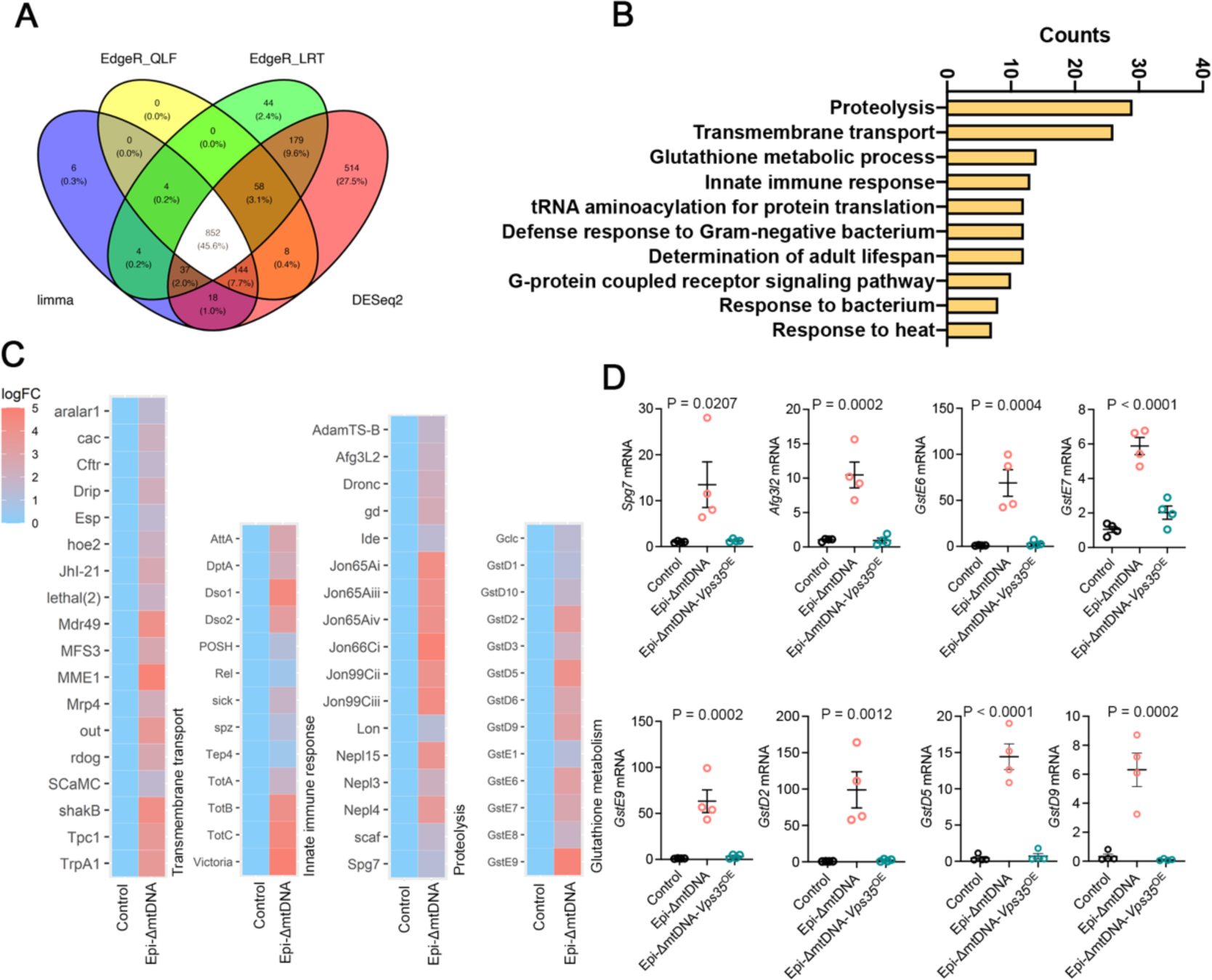
VPS35 expression restores mitochondrial defects associated to ΔmtDNA. (**A**) Venn diagram for genes differentially expressed (FDR of 5%) between control and Epi-ΔmtDNA shared through 4 differential expression methods (Limma-voom, EdgeR-QLF, EdgeR_LRT and DESeq2). (n=4 per genotype). (**B**) DAVID Pathway enrichment analysis for genes differentially expressed. (**C**) Heat map for log fold change showing gene identities for the top four categories of differentially enriched pathways. (**D**) mRNA quantification of selected genes in Epi-ΔmtDNA and Epi-ΔmtDNA-*Vps35*^OE^ larvae. (n=4 per genotype). *P* values were calculated using One-way ANOVA with Tukey correction for multiple comparison. Data is presented as mean ± SEM.

Epi-*Vps35^OE^* in Epi-ΔmtDNA larvae (Fig. S5C), reversed to normal levels all mRNA levels for the selected targets (Figure 8D). Thus, these data confirmed that *Vps35*^OE^ is sufficient to restore mitochondrial abnormalities caused by Epi-ΔmtDNA.

## Discussion

Mitochondrial quality control serves as a salvage pathway to eliminate ill-functioning mitochondria. The existence of alternative or cargo-specific mitophagy pathways highlights the complexity and adaptability of cellular systems. Different studies showed that these pathways share components with the vesicle trafficking system. For instance, the retromer, traditionally involved in endosomal maturation^27, 28^ induces, upon certain conditions, mitochondrial-derived vesicles containing oxidized cargo directed to peroxisomes^29^. Other proteins from the RAB family such as RAB9, RAB35, and RAB10, contribute to mitochondrial quality control and determine cargo selection^14, 30–33^. Recently, we found that persistent mtDNA replication stress does not trigger bulk mitochondria removal, but degradation of mitochondrial nucleoids^6^. Thus, these selective degradation pathways of damaged mitochondrial components ensure the elimination of focal damage, sparing degradation of the complete organelle and in the case of mitochondrial nucleoids, restricting the expansion of mtDNA mutations^34^.

The most common mechanism of mtDNA damage and mitochondrial dysfunction is the random accumulation of mtDNA mutations during aging^35^. Mutated mtDNA, encoding essential genes for mitochondrial function, leads to an imbalance of the respiratory chain and an increase in oxidative stress^36^. This enhances the mtDNA mutation rate by interfering with mtDNA processes, such as replication and transcription, posing a challenge to mitochondrial fitness. Previously, we found that persistent mtDNA replication stress leads to oxidative stress in mitochondrial nucleoids and translocation of the mtDNA to recycling organelles^6^. In agreement with previous data, through proximity labeling between mitochondria and endosomes, we detected a sum of proteins related to vesicle trafficking or lysosomal components. Among them, are the small GTPase RAB10 and the retromer component VPS29. RAB10, traditionally associated with vesicle transport and ER morphology^37^ also controls mitochondrial turnover upon depolarization^14^. Recently, RAB10 was found to be recruited in active lysosomes^38^. Noteworthy, RAB10 dynamics is dependent on LRRK2, a kinase involved in several steps of macro-autophagy activation^39^. Thus, by recruiting autophagy receptors and active lysosomes to the mitochondrial vicinity, RAB10 assists mitochondrial fission upon stress conditions and the removal of the leaked mitochondrial genome by lysosomes.

Delivery of mitochondrial components to recycling organelles has been shown to occur through mitochondrial-derived vesicles^16, 19, 40^. Sequential electron microscopy showed that at least the mtDNA is not expelled in a derived vesicle in the cytoplasm but is directly transferred to a degradation organelle. Whether mtDNA is expelled through a transient pore or selected through consecutive mitochondrial fission and engulfed in a lysosome, needs to be determined. Nevertheless, previous fragmentation of the mitochondrial network stimulated by VPS35 and RAB10 would protect the mitochondrial network by removing only ill-functioning parts. The fact that we can only detect matrix material, dsDNA or PDH, outside the mitochondrial compartment upon lysosomal inhibition, suggests that, leaked mitochondrial content is rapidly eliminated. Hence, our data supports a model through mitochondria-lysosome kiss-and-run, to eliminate the mitochondrial genome upon molecule damage.

Noteworthy, we did not observe the expel of mitochondrial matrix components in cells expressing the mutation VPS35^D620N^. This mutation in the core component of the retromer impairs lysosomal function^41^ and directly affects the delivery of the lysosomal enzyme cathepsin^18^. Interestingly, the retromer also participates in lysosomal development by promoting the maturation of hydrolases^42^. Other proteins detected in our proteomics approach such as ATP6V0D1, MYO1B, or BIRC6 are involved in lysosomal acidification and in the final steps of autophagosome-lysosome fusion^43–45^. Therefore, our data underlines the critical role of lysosomes in this pathway and suggests that healthy lysosomes are required to initiate the ejection of mitochondrial matrix material.

Given that the overexpression of VPS35 increases mtDNA extraction *in vitro*, we hypothesized that the modulation of the retromer *in vivo* would be beneficial to eliminate the burden associated with mtDNA damage. Noteworthy, overexpression of VPS35 has been shown to restore cellular defects associated with lysosomal dysfunction^46, 47^. In agreement, our data confirm that overexpression of *Vps35* in *Drosophila* can restore mitochondrial defects associated with altered mtDNA. The presence of ΔmtDNA rewires the mitochondrial redox state, which is translated into activation of detoxifying pathways. These changes activate a specific transcriptome profile directed to counteract cellular damage, but also to modify mitochondria structure. Thus, besides genes for glutathione metabolism and innate immune response, we detected upregulation of many transcripts for membrane carriers, such as members from aquaporin (*Drip*), ABCB transporters (*Mdr49*, *Mrp4*), and ion channels (*aralar1*, *MME1*, *sCaMC*, *TrpA1*). In addition, mitochondrial proteases such as *Lon*, *Afg3L2,* and *Spg7*, are all involved in the degradation of damaged mitochondrial proteins, and maintain mitochondria integrity^48^. Importantly, *Vps35*^OE^ leads to accelerated transfer of mitochondrial components to lysosomes, eliminating the source of cellular burden and modulation the downstream response.

SAMM50 and VPS35 reside at the crossroads of mitochondrial turnover pathways. Depletion of both proteins follows robust activation of mitophagy^6, 25^. In contrast, in oxidative conditions, while SAMM50 promotes piecemeal mitophagy of mitochondrial OXPHOS components^49^, overexpression of VPS35 enhances vesicle formation. Hence, both SAMM50 and VPS35 promote the activation of selective pathways and restrict the removal of the complete organelle. Paradoxically, the reduction of *Sam50* in the epidermis also restores the mtDNA copy number in Epi-ΔmtDNA larvae. In this case, a combination of mtDNA leak and a robust mitophagy response associated with the loss of SAM50 would eliminate ΔmtDNA generated in L2 larvae, inducing repopulation with wt mtDNA in mid-L3 larval instar, when the expression of transgenes is reduced.

Our study has been however directed into molecular aspects of mtDNA turnover, not considering physiological changes. We have not considered the effect of *Vps35* overexpression further than the *Drosophila* larval phase. Hence, at this stage, we cannot exclude, the pleiotropic effects caused by the overexpression, but undoubtedly, it initiates a new path. Further experiments are required to validate this approach in mammals.

In conclusion, we have provided a mechanistic view of a complex mechanism. Lysosomes carry out the elimination of mtDNA upon damage. RAB10 and the retromer facilitate mitochondrial fragmentation and lysosomal recruitment, which leads to mtDNA expulsion (Figure 9). Interestingly, we prove that *in vivo*, the retromer can drive the selection of mutated mtDNA and hence, exemplifies a valid strategy to counteract mitochondrial damage associated with ΔmtDNA.

**Figure 9.**
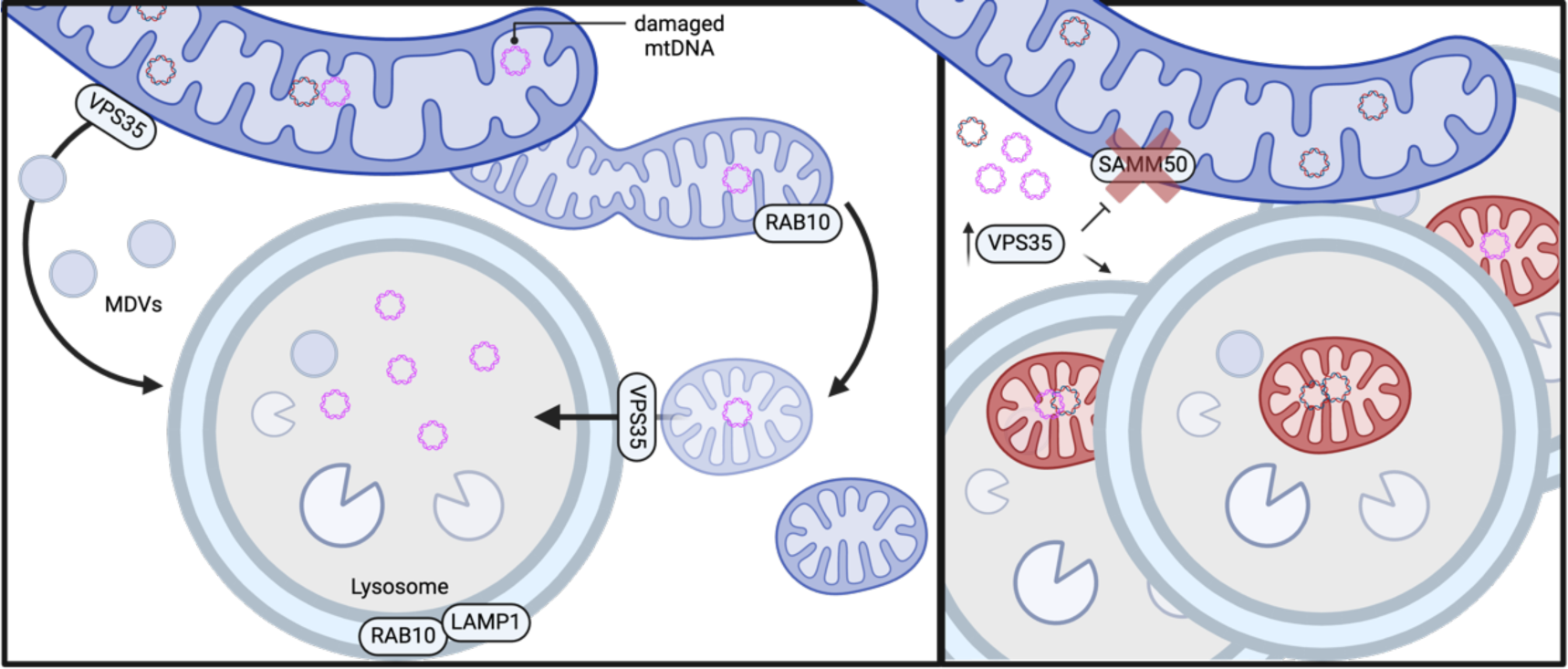
Proposed model. VPS35 enhances Mitochondrial derived vesicle and mtDNA release to lysosomes. RAB10 participated in mitochondrial fragmentation and lysosomal recruitment. Upon SAMM50 knock-down, overexpression of VPS35 enhances lysosomal recruitment and mitigates bulk mitophagy. Images created with biorender.com.

## Acknowledgements

We are thankful to the CECAD Imaging and Proteomics facility for excellent technical support, especially Christian Jungst for Light Imaging support, and Prerana Wagle for Proteomics analysis. We thank Berkan Akar and Jana Scholzen for technical assistance, Brue A. Hay, Bloomington, VDRC and DGGR Stock Centres for fly strains and Rudolf Wiesner and Maria Leptin for scientific discussion. This work was supported by grants from the Deutsche Forschungsgemeinschaft (PL 895/1-1) and Köln Fortune (186/2022) to D.P.M.

## Author contributions

Funding acquisition, D.P.M.; conceptualization, P.K, and D.P.M.; investigation and formal analysis, D.P.M., P.K., M.B., A.G., F.G., B.M., resources E.V., A.W., A.C.S; visualization, D.P.M.; writing-review & edit, P.K, and D.P.M.; Supervision, D.P.M.

## Material and methods

### Plasmid Generation

VPS35-EGFP and VPS35-CFP were generated by subcloning VPS35 from pLenti/V5/DEST-VPS35 plasmid (addgene #21691) in pEGFP-N1 and pECFP-N1 plasmid using XhoI and BamHI sites. Twinkle Homo sapiens ORF was subcloned from Twinkle-APEX2-V5 (addgene #129705) in pN1-mCherry using BglII and EcoRI sites. VPS35 p.D620N and Twinkle p.K319E were generated by site-directed mutagenesis. All plasmids were verified by Sanger sequencing. To generate stable cell lines, VPS35 or VPS35 p.D620N ORFs were subcloned in pLenti/V5/DEST using EcoRV sites

### Cell culture and chemical treatments

Cells were maintained in DMEM 4.5 g/L Glucose + GlutaMax (Gibco, #10566016) supplemented with 10% FBS, 1x Pen/Strep (Gibco, #15070063) and 2.5 μg/ml Plasmocin (Invivogen, #ant-mpp). Transfection of HeLa cells with VPS35 plasmids was achieved using FuGene HD (Promega, #E2311) following the manufacturer’s instructions. The lysosomal activity was inhibited with 10 μM Chloroquine (Sigma, #PHR1258) for 4h. Oxidative damage was induced by treating cells with 200 μM H_2_O_2_ in a complete medium for 4h. For stimulating mitochondrial metabolism, cells were grown overnight in non-glucose DMEM supplemented with 10mM Galactose (Sigma, #1287700), 10% FBS, 1x Pen/Strep, 1x GlutaMax (Gibco, #35050061) and 1x Pyruvate (Gibco, #11360070). SAMM50 Knock-Down (KD) conditional HeLa cells were kindly provided by Dr. Vera Kozjak-Pavlovic and described before^24^. *SAMM50* KD was induced by adding doxycycline 1 μg/ml (Sigma, #D3447). Medium was replaced every 48 hours and cells were used after 5 or 10 days.

For generation of stable cell lines, HeLa cells were transduced with pLenti/V5/DEST-VPS35 and pLenti/V5/DEST-VPS35 p.D620N using psPAX2 and pMD2.G as helper plasmids and HEK293 as packaging cell. The supernatant containing lentivirus was filtered through a 0.45 μm filter and supplemented with 10 μg/ml Polybrene (Sigma, #H9268). Transgenic expression of VPS35 was confirmed by western blot. Transduced clones were selected and maintained in a medium containing 20 μg/ml Blasticidin S (Invivogen, #ant-bl-05).

### *Drosophila* stocks

The GAL4/UAS system was used for ectopic expression^50^. All *Drosophila* experiments were carried out using *A58-GAL4* driver lines to express UAS-constructs in the epidermis of *Drosophila* larvae^51^. *A58-GAL4* is expressed in the epidermis from the first larval instar stage onward^52–54^. UAS-transgenes used in this work are: *UAS-mitoT4lig*, *UAS-mitoAflIII* (BL 84979), UAS-mitoAflIII, UAS-mitoT4lig/CyO (Kindly provided by Bruce A. Hay)^22^, *UAS-Vps35.HA*^55^ (BL67152), *UAS-Sam50-RNAi* (VDRC 106193 KK). All stocks were in a white-eyed genetic background and maintained at 25 °C under a 12:12 h light/dark cycle on standard fly food. All crosses were set up and maintained at 18°C under a 12:12 h light/dark cycle on standard fly food. All molecular experiments using *Drosophila* were conducted using complete larvae, excluding transcriptome analysis, where larval epidermis was isolated.

### mtDNA copy number and qPCR

Total DNA was isolated using the DNeasy Blood & Tissue Kit (Qiagen, #69504) according to the manufacturer’s instructions. 20 ng of total DNA was used for the analysis of threshold amplification differences between mtDNA and nuclear DNA (delta C(t) method. For human (HS) cell-based analysis, mtDNA copy number was obtained with oligos amplifying the mitochondrial gene *ND1* and the nuclear gene *APP*.

For cytosolic mtDNA detection, HeLa cells stably expressing VPS35 or D620N were transfected with Twinkle-K319E. 48h post-transfection, cells were collected and equally divided for internal control and fractionation experiments. The total DNA for internal control samples was obtained by solubilizing cell pellets in NaOH 10 mM, followed by boiling at 98°C for 30min and pH neutralization with Tris pH 8 to a final concentration of 150 mM. For mitochondria-free fractions, cell pellets were incubated in 25 μg/ml digitonin (Roth, #4005.2), 150 mM NaCl, and 50 mM HEPES pH 7.4 for 10 min on ice. Unbroken cells and mitochondria were cleared by sequential differential centrifugation first at 1.000 g, and resultant supernatant at 14.000 g. DNA was further isolated using the DNeasy Blood & Tissue Kit and eluted in equal volumes. 3 μl of pure DNA was used to amplify the nuclear gene *APP* and mitochondrial genes *ND1* and *ND5* in both fractions. Cytoplasmic content of mtDNA was obtained after normalization of each Ct value from mitochondria-free fractions to normalized value of total sample Ct (Ct Mitochondrial^MITOC.Free^ – (Ct Mitochondrial^TOTAL^ – Ct Nuclear^TOTAL^) = DDCt. Fold change values were obtained with 2^-DDCt.

For mtDNA heteroplasmy quantification and mtDNA copy number in *Drosophila*, 20 ng of total DNA isolated from 5 larvae was used to amplify *ND5* and *CytB*, mitochondrial genes outside and inside the deleted region respectively, and *3RTub* (TUBB) as a nuclear gene. Both Ct values from mitochondrial genes were normalized with the Ct for *TUBB* and the ratio between *CytB/ND5* was calculated. Quantitative real-time PCR for gene expression was performed using cDNA retrotranscribed from 1 μg of total RNA using PowerUP SYBR green (Thermo Scientific, #A25780).

All qPCR experiments were performed in Quant Studio 1 and SYBR Green from Thermo Scientific. Conventional PCR was performed using either cDNA or total DNA, specified in the corresponding figure. Primer sequences are specified in Table 1 for HeLa cell-related experiments and Table 2 for *Drosophila*.

**Table 1.**
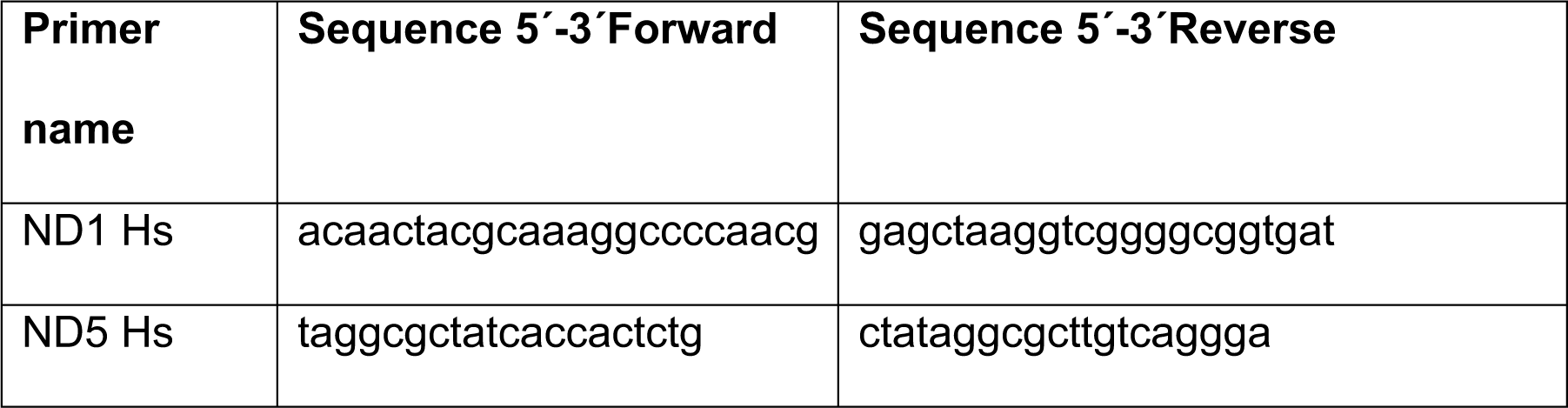
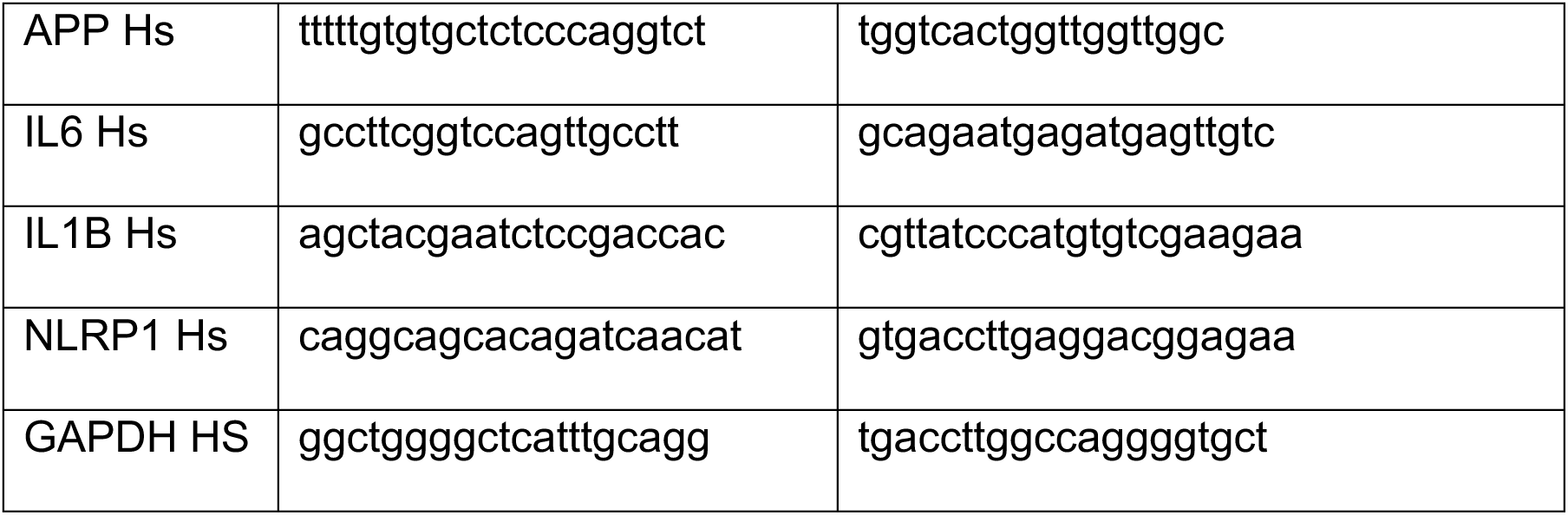
Primer sequence for HeLa cells-related analysis.

**Table 2.**
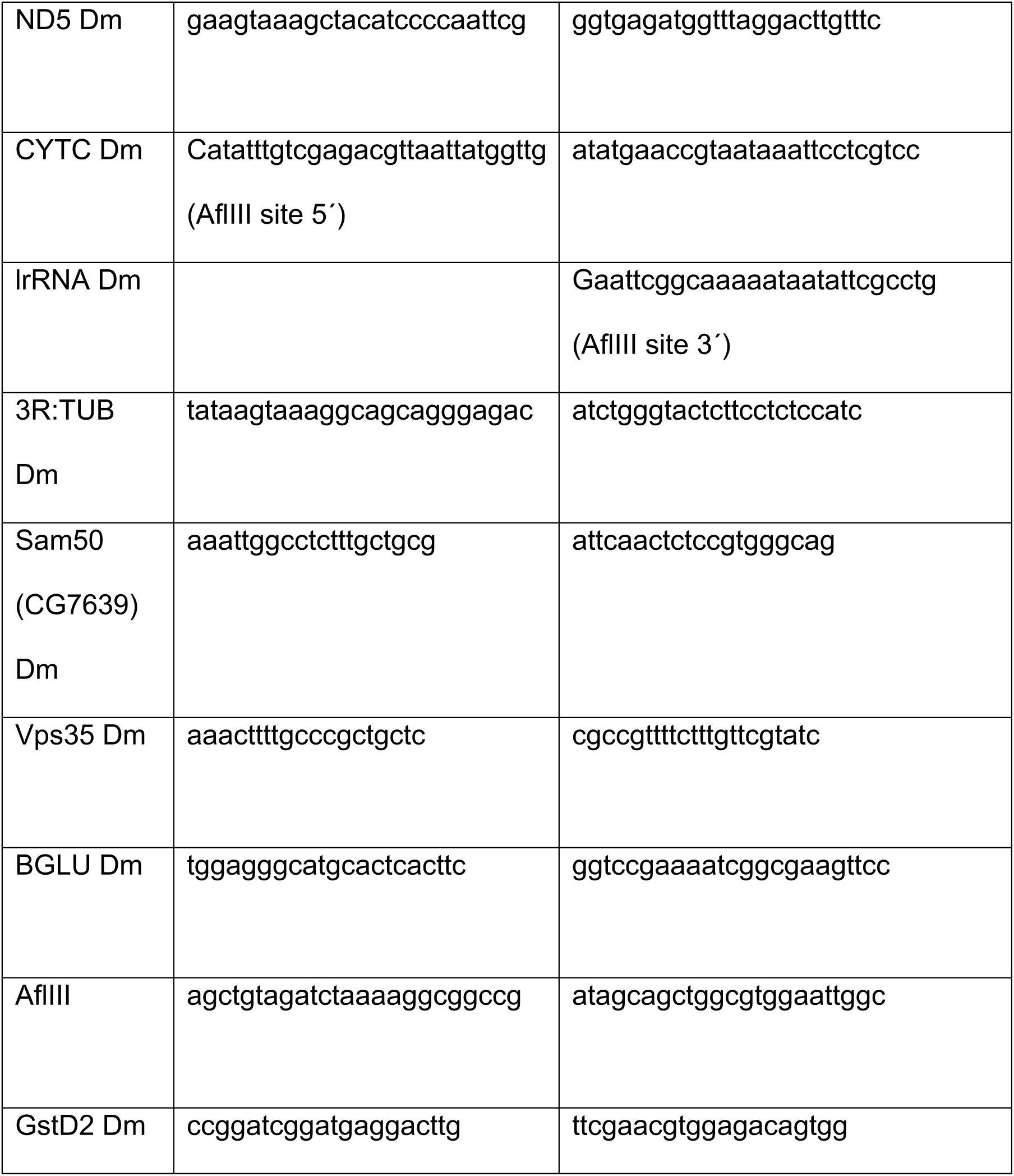
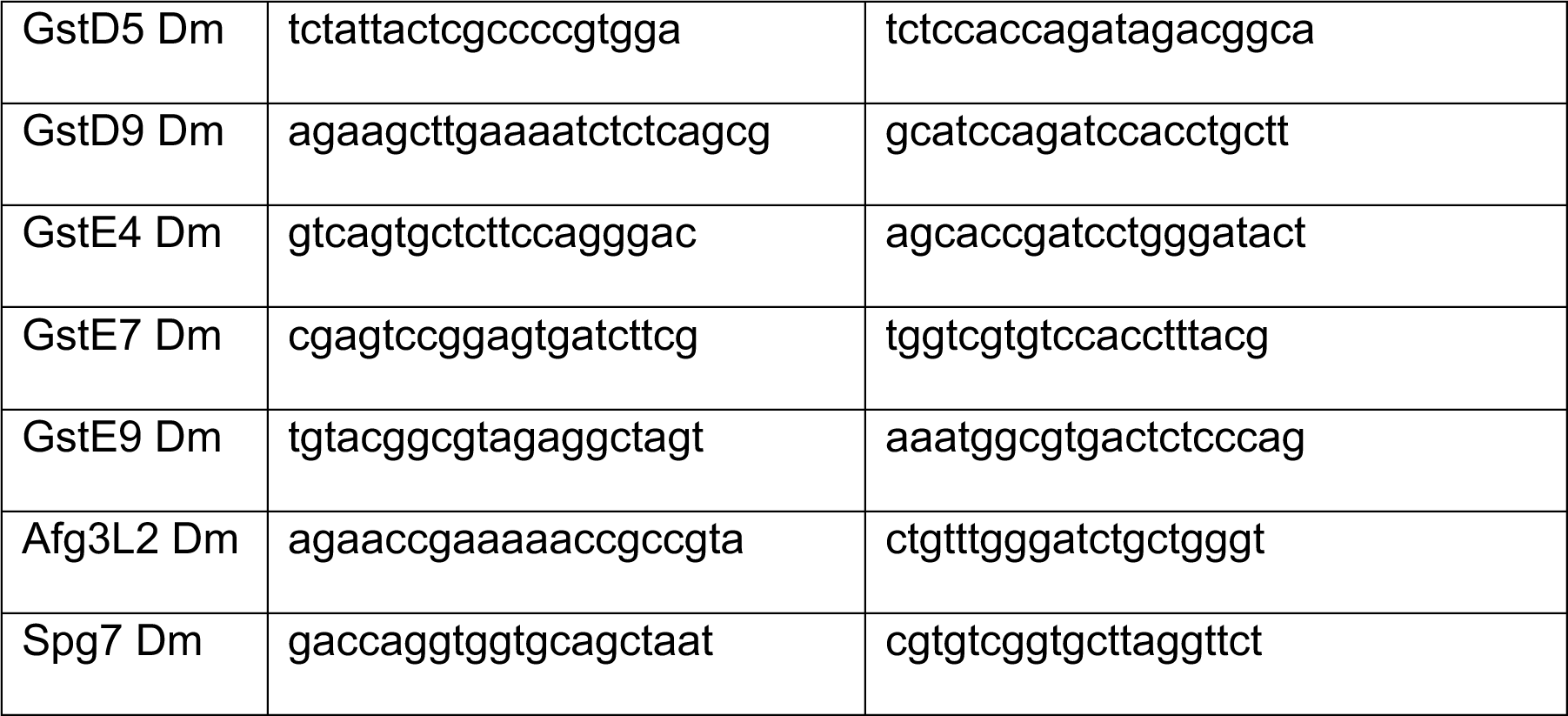
Primer sequence for *Drosophila-*related samples.

### Western blot and immunofluorescence

For protein isolation, cells were pelleted at the indicated time points and resuspended in RIPA Buffer (150 mM NaCl, 1% Triton-X100, 0.1% SDS, 50 mM Tris-HCL pH 8, 1 mM EDTA, 0.5% Na-deoxycholate) containing protease inhibitor (Sigma, #11697498001). Protein concentration was quantified using Bradford reagent (Bio-Rad, # 5000001), and equal amounts of protein were loaded in SDS-PAGE, transferred to PVDF membrane (XYZ), blocked in TBST-5% Fat-free milk, and blotted with the indicated antibodies. Antibodies used for western blot were: polyclonal α-SDHB (10620-1-AP; 1:1000), α-optineurin (10837-1-AP; 1:1000) α-LC3B (14600-1-AP; 1:1000), α-GFP (50430-2-AP; 1:1000) and monoclonal α-GAPDH (60004-1-Ig; 1:4000), α-GFP (66002-1-Ig; 1:1000) and α-p62 (66184-1-Ig; 1:1000) from Proteintech; polyclonal α-SAMM50 (ab133709; 1:1000), α-RAB5 (ab109534; 1:1000) and monoclonal α-ATP5A (ab14748; 1:1000) from Abcam; polyclonal α-LAMP1 (#9091; 1:1000), α-V5 (#13202; 1:1000) and monoclonal α-V5 (#80076; 1:1000) from Cell Signaling; monoclonal α-VPS35 (sc-374372; 1:1000) and α-RAB5 (sc-46692; 1:500) from Santa Cruz. For western blot involving *Drosophila*, 5 L3 larvae were homogenized in lysis buffer (150 mM NaCl, 1% Triton-X100, 50mM Tris-HCL pH 8, 0.01% NP-40, 1mM EDTA) containing a protease inhibitor, using a Teflon pestle. Equal amounts of proteins were directly loaded in SDS-PAGE and proceeded as described. All membranes were incubated with appropriate secondary antibodies combined with HRP (Jackson ImmunoResearch; goat anti-mouse #115-035-003; goat anti-rabbit #111-035-003), developed using Western Lighting Plus ECL (PerkinElmer, #NEL104001EA), and visualized using LAS500 CCD camera (GE Healthcare) For immunofluorescence, cells were seeded on glass coverslips and fixed for 15 min in 4% PFA/PBS and stored at 4°C until use. Permeabilization was achieved using 0.2% Triton/PBS followed by 1h blocking (5% Fat-free milk, 10% FBS, 0.1% Triton, 0.5% BSA in PBS). Primary antibodies were incubated overnight at 4°C. Antibodies used for immunofluorescence were: polyclonal α-TOM20 (11802-1-AP; 1:1000), α-Calnexin (10427-2-AP; 1:500) from Proteintech; monoclonal α-dsDNA (ab27156; 1:1000), α-PDHX (ab110333; 1:500), α-ATP5A (ab14748; 1:500) and polyclonal α-RAB5 (1:500) from Abcam; polyclonal α-LAMP1 (#9091; 1:500), α-V5 (#13202; 1:500) and monoclonal α-V5 (#80076; 1:500) from Cell Signaling; monoclonal α-VPS35 (sc-374372; 1:500), from Santa Cruz. After washing the coverslips with PBS, samples were incubated with appropriate secondary antibodies (1:1000, 1h at room temperature) from Thermo Scientific: goat α-rabbit combined with Alexa-405 (#A-31556), goat α-rabbit Alexa-488 (#A-11008), goat α-mouse Alexa-488 (#A-11001), goat α-rabbit Alexa-546 (#A-11035), goat α-mouse Alexa-546 (#A-11030), goat α-rabbit Alexa-647 (#A-21245) and goat α-mouse Alexa-647 (#A-21235) Coverslips were mounted using Fluoromount containing DAPI (Thermo, #00-4959-52).

### Confocal and Super-resolution microscopy

For confocal microscopy, Images were acquired in Leica SP8 with 63x/1.40 oil PL Apo-objective, Zeiss Airy Scan with 63x/1.40 oil DIC, and PerkinElmer Spinning-disk confocal microscope UltraVIEW VoX with 60x objective. All images were deconvoluted using ImageJ (NIH) before analysis to increase resolution. Brightness was adjusted equally in the entire image.

For Super Resolution of cristae structure, cells were plated in glass bottom plates (Ibidi, #81158) and loaded 1h with 250nM PK MitoOrange (Spirochrome, # SC053). Images were acquired using TCP SP8 gSTED Leica Microsystems using PL Apo x100/1.40 Oil STED Orange. Airy Scan microscopy was used also to determine cytosolic mtDNA in HeLa cells upon *Samm50* KD.

All image analyses were performed using ImageJ (NIH). Quantification of mitochondrial morphology was achieved using a Mitochondrial morphology macro^56^. MDV quantification was performed using an internal plugin by counting TOM^-^ PDH^+^ particles. Manderś coefficient was calculated using the JACOP plugin^57^. The fluorescence profile was obtained by drawing a line covering 50 pixels and using the RGB Profiler plugin.

### Electron microscopy

Epidermis from *Drosophila* L3 larvae were isolated and emersion fixed in 2% formaldehyde, 2% glutaraldehyde, and 3 mM CaCl_2_ in 0.1 M sodium cacodylate buffer for 48 h. Samples were washed with 0.1 M sodium cacodylate buffer and postfixed with 2% OsO_4_ in 0,1M cacodylate buffer, washed 3×5 min with ddH2O, followed by dehydration in an ascending ethanol series for 15 min each (50 %, 70 %, 90 %, 3×100 %) at 4°C. Samples were incubated for 15 min with 50% ethanol/propylene oxide, followed by two times 15 min pure propylene oxide. Then, samples were infiltrated with a mixture of 50% epon/propylene oxide and 75% epon/propylene oxide for 2 h each, and pure epon overnight at 4°C. The next day, samples were incubated with fresh epon for 2h and, placed into flat embedding moulds and cured at 60°C.

Ultrathin sections of 70 nm were cut using an ultramicrotome (Leica Microsystems, UC6) and a diamond knife (Diatome, Biel, Switzerland) and stained with 1.5 % uranyl acetate and 3 % Reynolds lead citrate solution. Images were acquired using a JEM-2100 Plus Transmission Electron Microscope (JEOL) operating at 80kV equipped with a OneView 4K camera (Gatan).

For CLEM, cells were seeded onto carbon-coated plates containing a numerated grid. Cells were transfected with TWNK^K319E^-Cherry and VPS35-CFP plasmids as described previously. The lysosomal function was blocked for 4h with 10 μM chloroquine. 1h before fixation, cells were loaded with 1x SYBRGold (Thermo, # S11494) and 250nM PK mito Deep Red (Spirochrome, # SC055). Cells were fixed for 30 min at room temperature and 30 min at 4°C in 2 % glutaraldehyde, 2.5 % sucrose, 100mM CaCl_2_ in 0.1M HEPES pH 7.4 and washed with 0.1M HEPES buffer. Transfected cells showing cytoplasmic DNA were scanned using the SP8 confocal microscope (Leica) with 63x/1.40 oil objective and brightfield images were used to localize the coordinates of cells of interest. Following light imaging, cells were incubated with 1% Osmiumtetroxid for 30 min at 4°C, washed in 0.1M Cacodylate buffer, and dehydrated using ascending ethanol series (50 %, 70 %, 90 %, 100 %) at 4°C. Cells were infiltrated with a mixture of 50% Epon/ethanol for 1 h, 66% Epon/ethanol for 2 h, and pure Epon overnight at 4°C. TAAB capsules filled with Epon were placed upside down onto the glass bottom and cured for 48 h at 60°C. Glass bottom was removed by alternatingly putting the dish into boiling water and liquid nitrogen. Block face was trimmed to the previously noted coordinates using a 90° trim tool (Diatome, Biel, Switzerland) and ultrathin sections of 70 nm were cut using an ultramicrotome (Leica Microsystems, UC6) and a diamond knife (Diatome, Biel, Switzerland), stained with 1.5 % uranyl acetate for 15 min at 37°C and lead citrate solution for 4 min. For electron tomography, Ultrathin sections of 300 nm were cut and incubated with 10 nm protein A gold (CMC, Utrecht) diluted 1:60 in ddH20. Sections were stained with 2% Uranyl acetate for 20 min and Reynolds lead citrate solution for 3 min. Images for the Tilt series were acquired with a pixel size of 1.108 nm from −60° to 60° with a 1° increment on the same JEM-2100 Plus Transmission Electron Microscope (JEOL) operating at 200 kV. Reconstruction was done using Imod, MiB and Imaris.

### Proximity biotinylation

Proximity biotinylation was performed in transduced cells expressing Split TurboID constructs RAB5C-SplitN-V5 and SAMM50-SplitC-HA. Cells expressing only SAMM50-SplitC-HA were used as a negative control.

For protein isolation, HEK293 cells were incubated for 4 h with 0,25 mM biotin. For inducing mtDNA damage, cells were previously transfected with Twinkle K319E-Cherry 24 h before the biotin incubation. Pellets were scraped from plates right after the biotin incubation, and solubilized in RIPA buffer. 300 µg of total protein extracts were purified using Streptavidin Sepharose High-Performance beads (GE Healthcare Bio-Sciences AB). For western blot analysis, beads were washed 3 times with RIPA buffer, resuspended in 2x Laemmli buffer, boiled at 95°C for 5 min and loaded onto SDS-PAGE. Biotinylated proteins were visualized with α-Streptavidin-HRP antibody (Jackson ImmunoResearch, 1:2000. #016-030-084). For mass spectrometry, beads were washed 3 times with ABC buffer (Ammonium Bicarbonate 50 mM pH 7.8), incubated 10 min with 50 µl of Urea buffer (6 M urea, 2 M thiourea), 1h with 5 mM dithiothreitol (DTT) and 30 min with 40 mM Indol-3-acetic acid (IAA) at room temperature and in the dark. Protein digestion was achieved by incubating with Lys-C in a ratio of 1:100, for 2-3 h. Samples were diluted with ABC buffer to a final urea concentration of 2 M, and incubated with Trypsin overnight in a ratio of 1:100. Digestion was stopped with 1% formic acid and peptides loaded into pre-equilibrated stage tips and stored at 4°C until used for mass spectrometry analysis.

### Liquid chromatography-mass spectrometric analysis

Affinity-enriched proteomics samples were analyzed in positive mode using data-dependent acquisition (DDA) either by an Easy-nLC 1000 – Q Exactive Plus or an Easy-nLC 1200 – Orbitrap Eclipse tribrid system (Thermo Fisher). On-line chromatography was directly coupled to the mass spectrometric systems using a nanoelectrospray ionization source. Peptides were separated by reversed-phase chromatography with a binary buffer system of buffer A (0.1% formic acid in water) and buffer B (0.1% formic acid in 80% acetonitrile) using a 60-minute chromatographic gradient. Separation was performed on a 50 cm long in-house packed analytical column filled with 1.9 μM C18-AQ Reprosil Pur beads (Dr. Maisch). Using the 60 min chromatographic gradient peptide separation based on their hydrophobicity was performed by linearly increasing the amount of buffer B from initial 13% to 48% over 35 min followed by an increase of B to 95% for 10 min. The column was washed for 5 min and initial column conditions were achieved by equilibrating the column for 10 min at 7% B. Full MS spectra (300 to 1750 m/z) were acquired with a resolution of 70.000, a maximum injection time of 20 ms and an AGC target of 3e6. The top 10 most abundant peptide ions were isolated (1.8 m/z isolation windows) for subsequent HCD fragmentation (NCE = 28) and MS/MS recorded at a resolution of 35,000, a maximum injection time of 120 ms and an AGC target of 5e5. Peptide ions selected for fragmentation were dynamically excluded for 20 seconds.

### Data processing and analysis for proteomics

All recorded RAW files were processed with the MaxQuant software suite 75 (1.5.3.8 for IP data, 1.6.14 for pSILAC). For peptide identification and scoring MS/MS spectra were matched against the mouse Uniprot database (downloaded 08/15/2019) using the Andromeda search algorithm 76. For the affinity-enriched samples, multiplicity was set to one and trypsin/P was selected as digestive enzyme. Carbamidomethylation was set as a fixed modification and methionine oxidation or N-terminal acetylation was selected as a variable modification.

Peptides were identified with a minimum amino acid length of seven and a false-discovery rate (FDR) cut-off of 1% on the peptide level. Proteins were identified with FDR < 1% using unique and razor peptides for quantification. Label-free quantification was performed using the standard settings of the maxLFQ algorithm. Match between runs was activated.

Statistical analysis and visualization were done with the Perseus (1.6.5) and InstantClue software suits 77,78. LFQ intensities of IP samples were log2-transformed and filtered for proteins identified in at least two replicates of one condition. Missing values were imputed with the Perseus plugin ImputeLCMD using a deterministic minimal value approach (MinDet, q-value = 0.001) to simulate the lower detection limits of the mass spectrometer. To evaluate the principal components responsible for the variances between samples, we performed a principal component analysis. Further, we performed a two-sided Student’s t-test to identify significantly regulated proteins (S0 = 0.1, permutation-based FDR = 0.05, 500 randomizations). If not otherwise indicated, significantly enriched proteins were determined by a combination of FDR-corrected p-values and log2 protein fold changes (q-value < 0.05 and absolute log2 FC > 1). Pathway analysis was performed using Metascape^58^.

### Transcriptomics analysis

For RNA next-generation sequencing, L3 larval epidermis from 5 larvae was first isolated and pooled. Total RNA was extracted as explained previously and 2ug of RNA was sequenced using Poly A + selection. A total of 25-30 million reads per sample were obtained using paired-end 100 pb read length (Illumina NovaSeq 6000). A quality control analysis of the RNA-seq raw data was performed with FastQC v0.12.0 (Babraham Bioinformatics) and MultiQC v1.18 software^59^. 3’ poly(A) tail and the 3’ adapter from Illumina were trimmed with the TrimGalore 0.06.10 tool (Babraham Bioinformatics) to reach a PhredScore greater than 35 and adapter content lower than 1%. Trimmed reads were then mapped onto the Drosophila Melanogaster genome (Drosophila BDGPG.32, ENSEMBL) by using the RNA-seq aligner STAR 2.7.10b^60^, followed by a quality assessment of the alignment with Qualimap v2.2.1^61^. To estimate gene expression levels, mapped reads for all transcript variants of a gene (gene counts data) were counted with RSEM v1.3.3 software.

Quality control, trimming, mapping and counting steps were performed in an Ubuntu 22.04.2 LTS environment. Further analysis was run through the RStudio 2023.03.0 +386 (R version 4.2.2) software. Thus, gene counts were annotated with the AnnotationHub Bioconductor package and four differential expressions (DE) analyses were performed both by applying the negative binomial distribution with EdgeR-LRT (likelihood-ratio test), EdgeR-QL (quasi-likelihood)^62^ and DESeq2^63^ bioconductor packages or by applying empirical Bayes model using Limma-Voom analysis^64, 65^. Genes with a common differential expression among the four methods (cpm < 0.5, p < 0.05) were selected for the Gene Set Enrichment Analysis (GSEA) with Gene Ontology (GO) pathways by using the clusterProfiler Bioconductor package^66^ and DAVID Knowledgebase (NIH).

### Data availability

The mass spectrometry proteomics data have been deposited to the ProteomeXchange Consortium (http://proteomecentral. proteomexchange.org) via the PRIDE partner repository^67^ under the accession PXD049081. New Generation Sequencing data has been deposited under GEOarchive.

---

**Figure S1.**
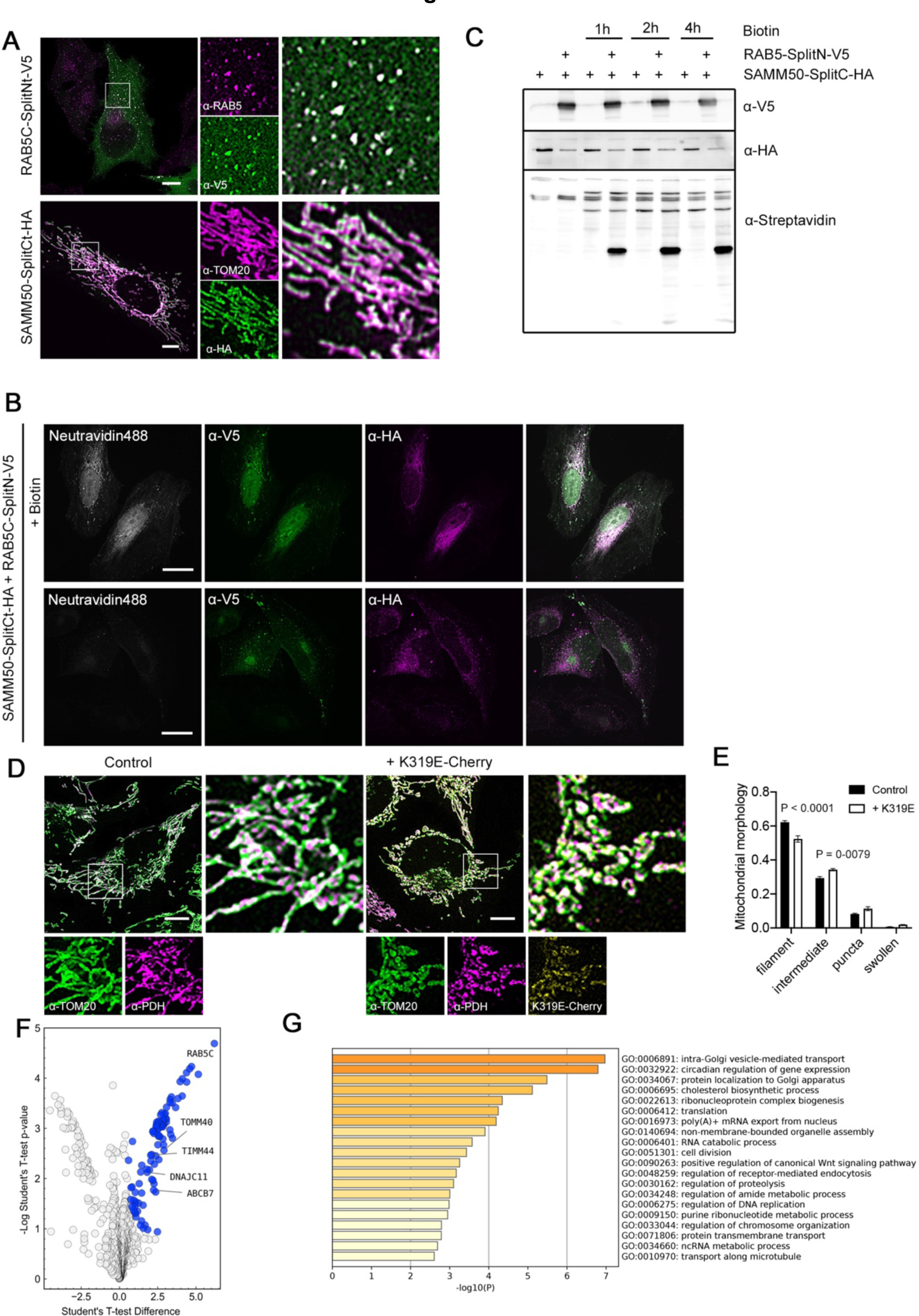
(**A**) Immunostaining of cells expressing RAB5C-SplitN-V5 and SAMM50-SplitC-HA showing specific localization in endosomes and mitochondria, by co-staining with α-RAB5 and α-HA, and α-TOM20 and α-HA respectively. (**B)** Reconstitution of biotinylation activity of TurboID by co-expression of RAB5-SplitN-V5 and SAMM50-SplitC-HA. Biotinylated proteins are detected with Neutravidin 488. α-HA and α-V5 were used to label SAMM50 and RAB5C. (**C**) Western blot analysis of biotinylated proteins labelled with α-Streptavidin-HRP. (**D, E**) Mitochondria morphology analysis in cells transfected with TWNK^K319E^-Cherry and labelled with the mitochondrial outer membrane marker α-TOM20 and mitochondrial matrix α-PDH. (n=3, >30 cells per replicate). (**F**) Volcano Plot showing proteins enriched after biotinylation and purification of cells transduced with SplitTurboID plasmids. Differentially expressed proteins compared with cells transduced with SAMM50-SplitCt (significant: q-value < -0.05 and absolute log2 fold change>1) are highlighted in blue. (n=3). (**G**) Pathway enrichment analysis with Metascape showing GO terms for proteins differentially enriched proteins. *P* values were calculated using Two-way ANOVA for genotype and morphology. Scale bar, 10 μm. Data is presented as mean ± SEM.

**Figure S2.**
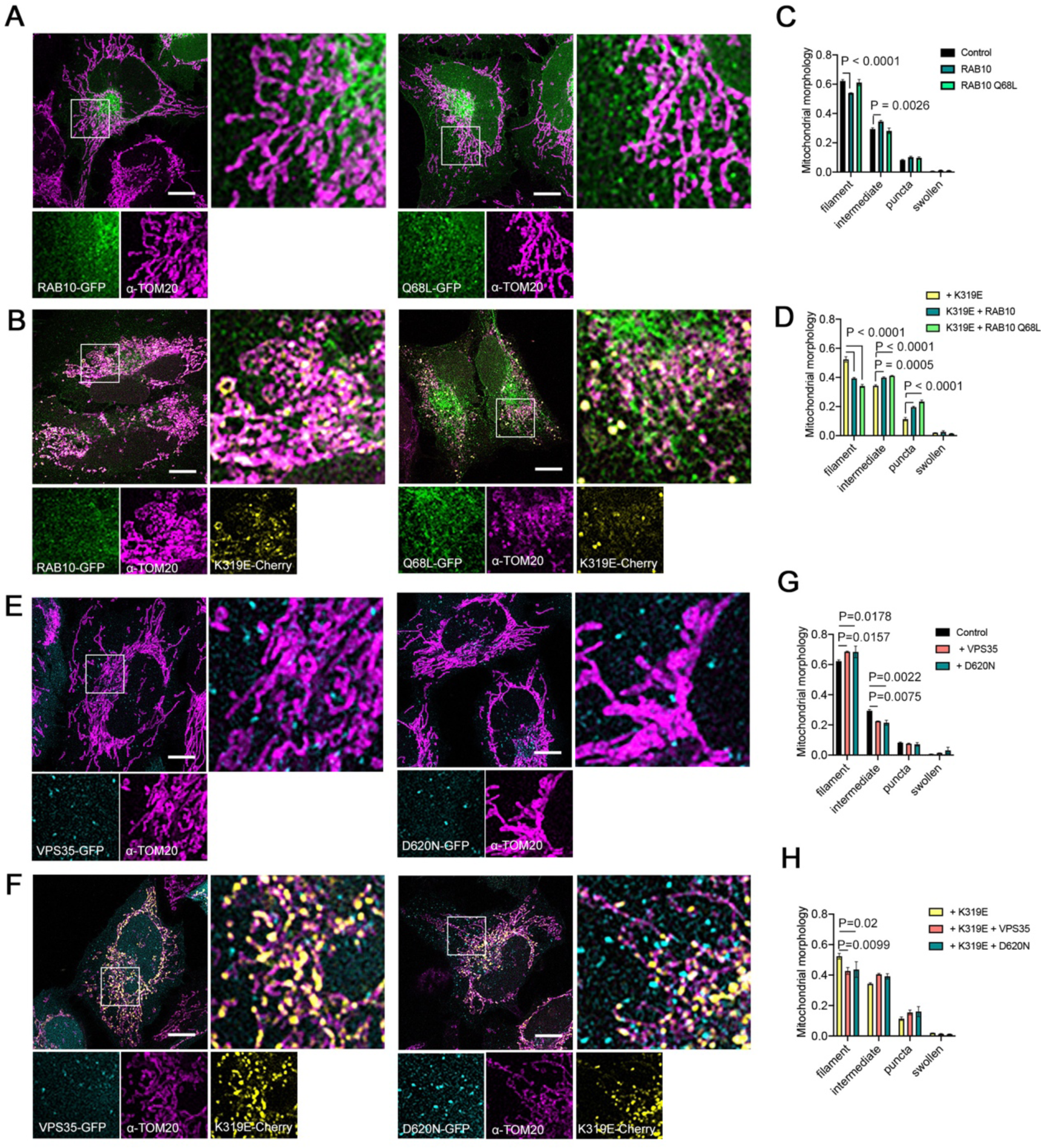
(**A**) Mitochondrial morphology analysis and quantification in cells transfected with RAB10-GFP, (**B**) RAB10^Q68L-^GFP, (**E**) VPS35-GFP, (**F**) VPS35^D620N^-GFP and in combination with TWNK^K319E^-Cherry. Mitochondrial morphology was obtained by analyzing staining with α-TOM20. (n=3, >30 cells per replicate). *P* values were calculated using Two-way ANOVA for genotype and morphology. Scale bar, 10 μm. Data is presented as mean ± SEM.

**Figure S3.**
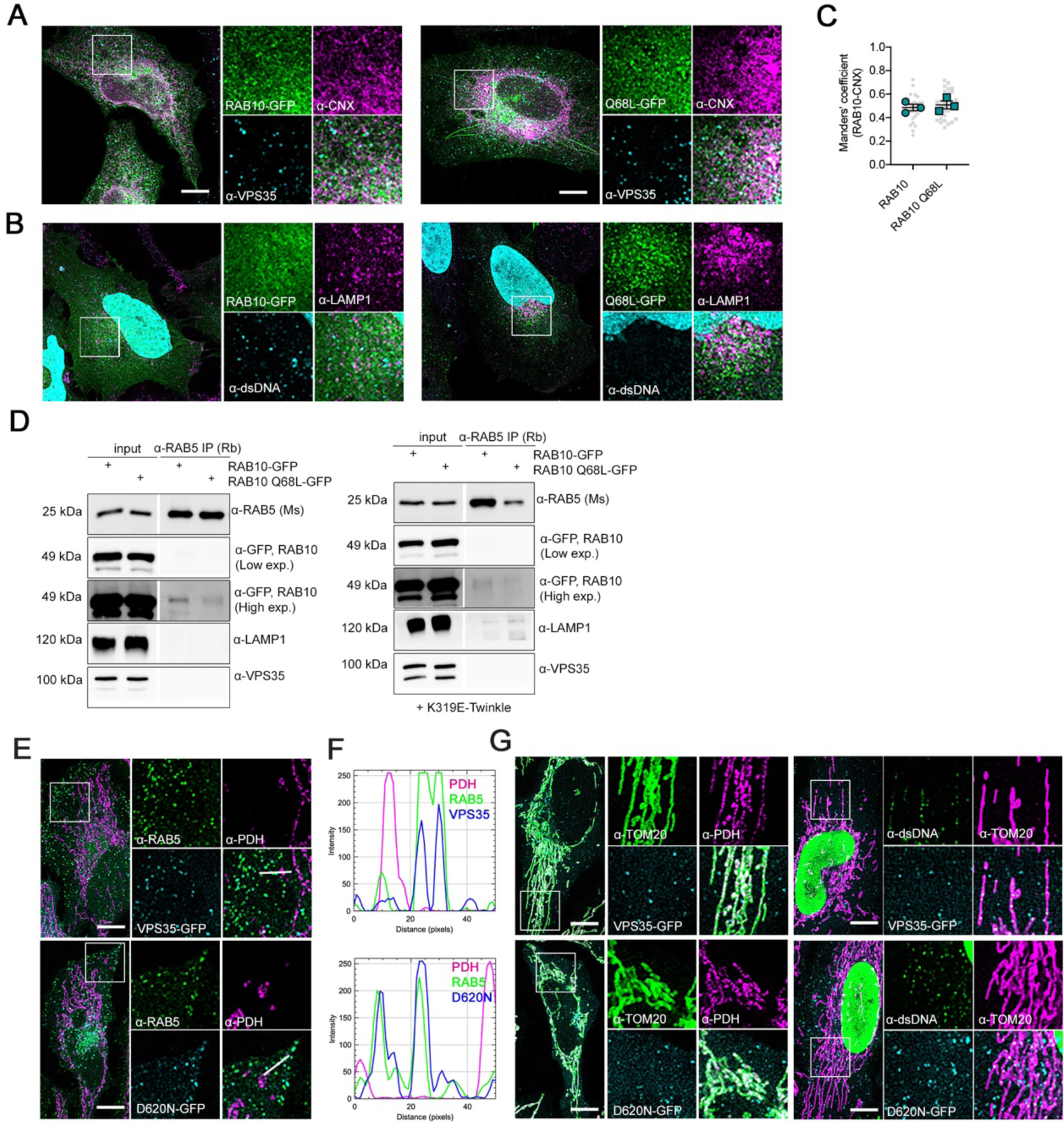
(**A**) Immunostaining of RAB10-GFP and RAB10^Q68L^-GFP transfected cells labelled with α-VPS35, the endoplasmic reticulum (ER) marker α-Calnexin (CNX) and (**B**) α-LAMP1 and α-dsDNA. (**C**) Manderścorrelation coefficient between RAB10 and Calnexin. (n=3, 10 images per replicate). (**D**) co-Immunoprecipitation of α-RAB5 with RAB10 and LAMP1. (**E, F**) Immunostaining and fluorescence profile of cells transfected with VPS35-GFP and VPS35^D620N^-GFP and labelled with α-RAB5 and α-PDH. (**G**) Cells transfected with VPS35-GFP plasmids and stained with mitochondrial markers α-TOM20, α-PDH and α-dsDNA. Scale bar, 10 μm. Data is presented as mean ± SEM.

**Figure S4.**
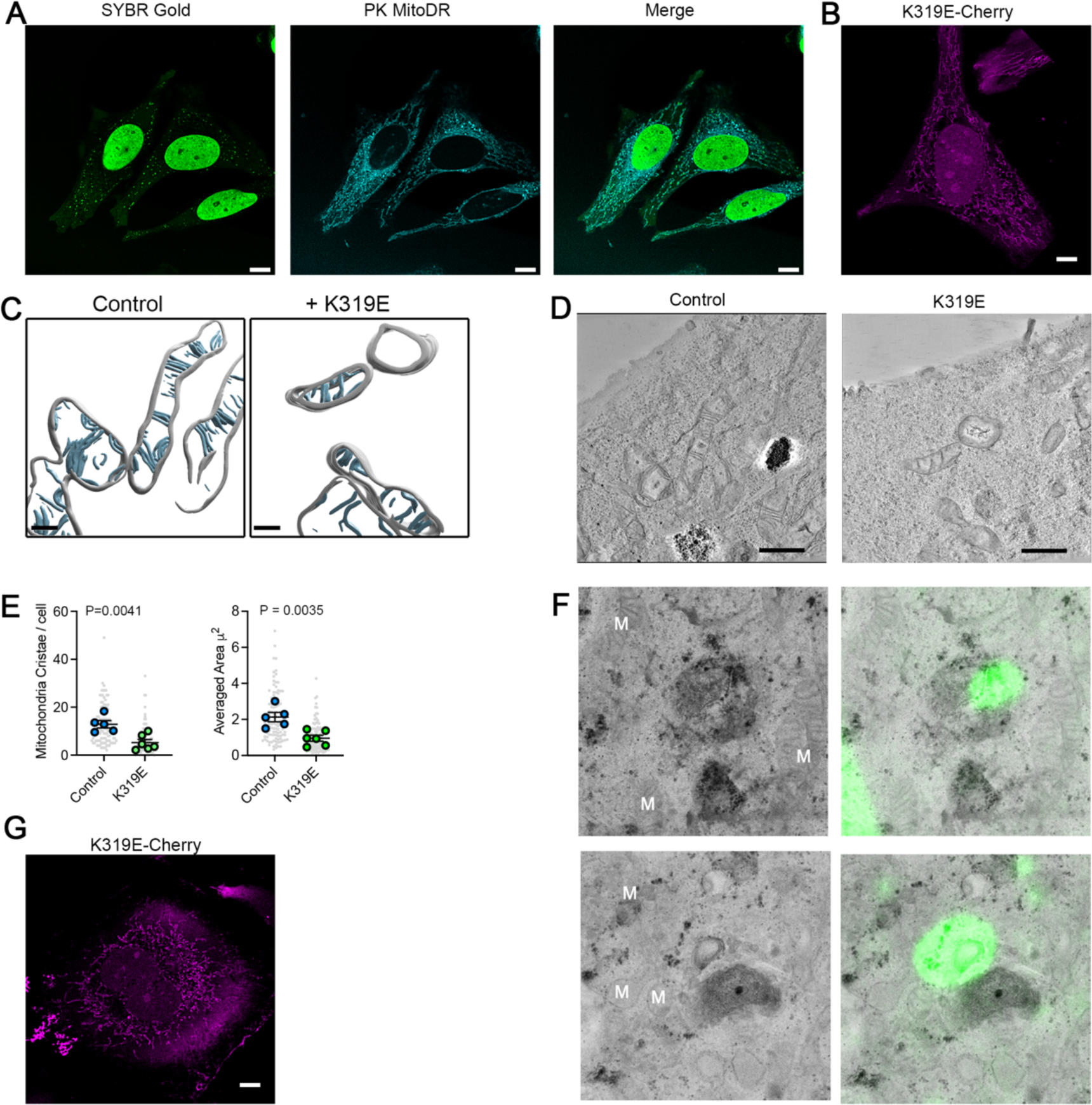
(**A**) Control cells loaded with SYBR Gold and PK MitoDeep Red. (**B**) TWNK^K319E^-Cherry corresponding to the cell used for CLEM in Figure 4A. (**C, D**) Volumetric reconstitution of electron tomographies from control and TWNK^K319E^-Cherry cells. (**E**) Quantification of the morphological parameters of mitochondria cristae based on electron microscopy images. (Control, n=5; TWNK^K319E^, n=6. >10 mitochondria per cell). (**F**) CLEM of cytosolic DNA presented in Figure 4A, B and C. (**G**) TWNK^K319E^-Cherry for the cell used for CLEM in Figure 4D. *P* values were calculated using Studentś T-test. Scale bar, 10 μm (A, B and G), 300 nm (**C**) and 1 μm (**D**). Data is presented as mean ± SEM.

**Figure S5.**
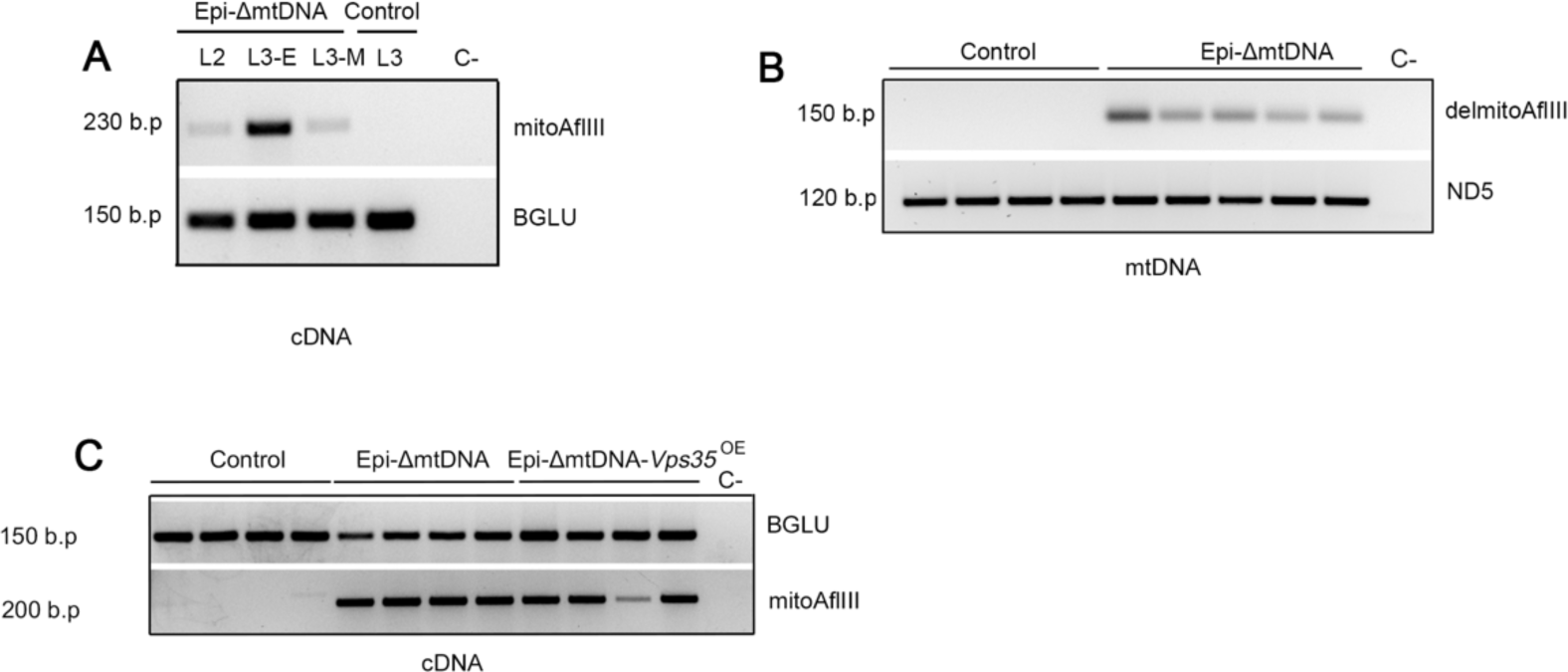
(**A**) Conventional PCR amplification of mtAflIII restriction enzyme in cDNA obtained from RNA isolates of L2 and L3 larvae (E, early; M, mid stage). mRNA for *BGLU* was used as housekeeping gene. (**B**) PCR amplification of the mtDNA deletion generated by co-expression of mitoAflIII and mitoT4-Ligase. Primers flank a region separated approx. by 2.500 b.p. Upon restriction of the mtDNA and ligation, the region is reduced to approx. 150 b.p. The mtDNA gene *ND5,* located outside the restricted region, was used as a control for mtDNA amplification. (**C**) PCR amplification of mitAflIII mRNA in Epi-ΔmtDNA and Epi-ΔmtDNA-*Vps35*^OE^ larvae. mRNA for *BGLU* was used as housekeeping gene.

**Figure S6.**
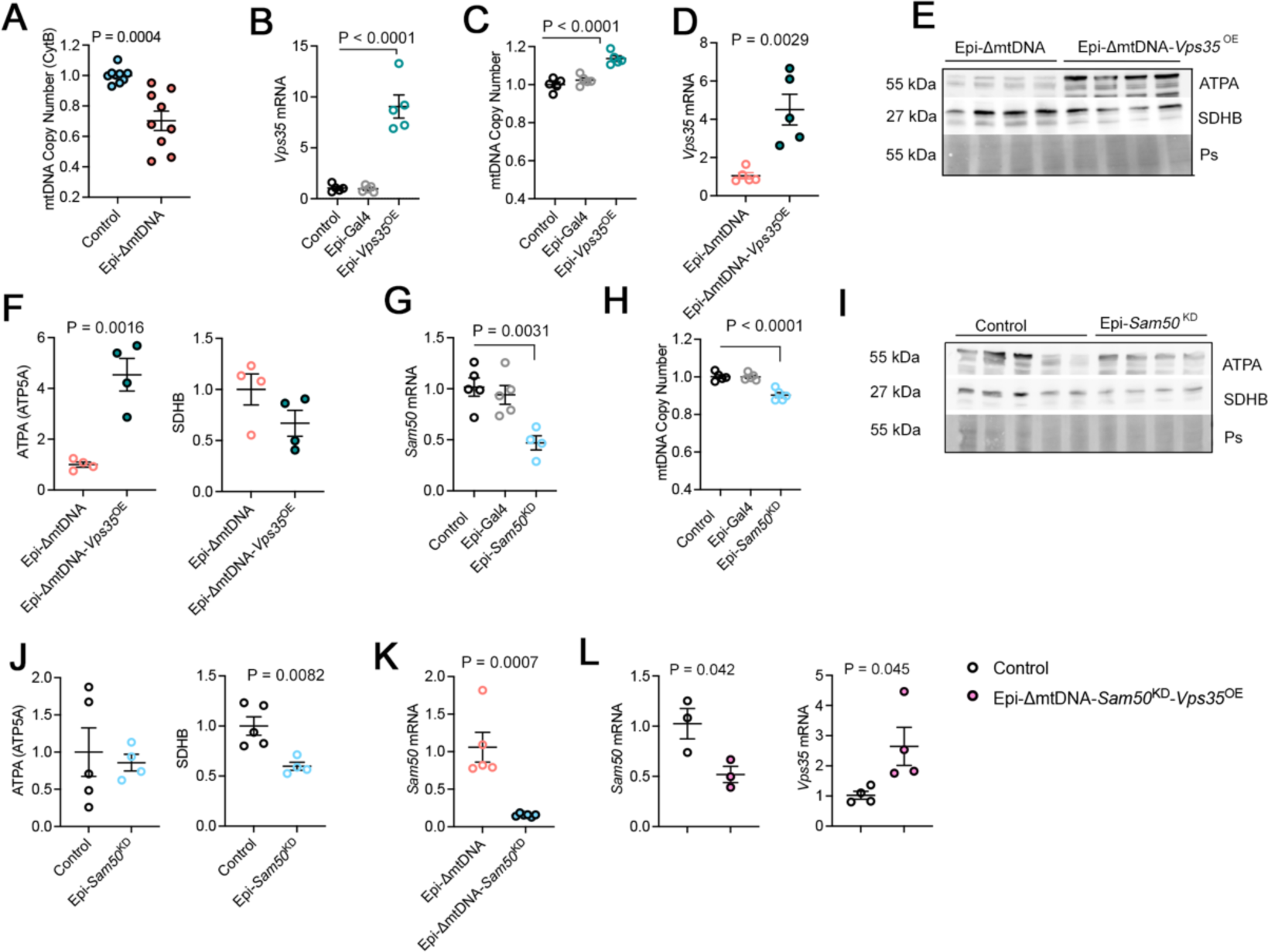
(**A**) mtDNA copy number analysis in Epi-ΔmtDNA larvae using *CytB* gene as a reference. (n=9). (**B, C, D**) mRNA quantification and mtDNA copy number analysis in Epi-*Vps35*^OE^. (n=5). (**E, F**) Western blot and quantification from total protein extracts for ATPA and SDHB for Epi-*Vps35*^OE^. Ponceau S (Ps) was used as a loading control. (n=4). (**G**) mRNA quantification and (**H**) mtDNA copy number in Epi-*Sam50*^KD^. (n=5). (**I**) Western blot and quantification from total protein for Epi-*Sam50*^KD^. Ponceau S (Ps) was used as a loading control. (Control, n=5; Epi-*Sam50*^KD^, n=4). (**K, J**) mRNA quantification for indicated genes in the selected genotypes. (n=3-6). *P* values were calculated using One-way ANOVA with Tukey correction for multiple comparison (B, C, G, H) and StudentśT-test (A, D, F, J, K, L). Data is presented as mean ± SEM.

**Figure S7.**
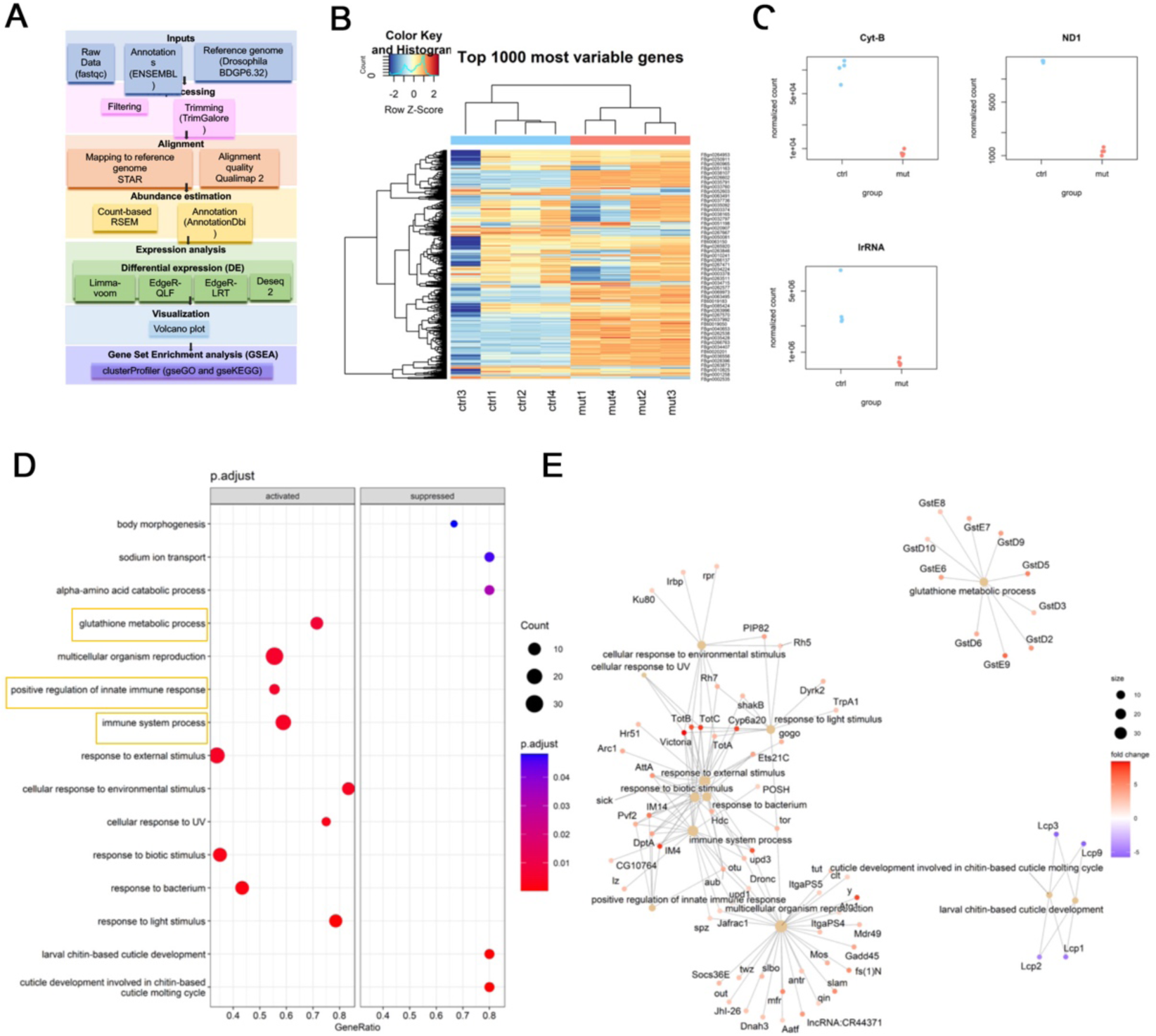
(**A**) Next Generation Sequencing (NGS) analysis flow chart for sequential steps and bioinformatic methods for the raw gene expression quantification in Ubuntu, following by the differential gene expression quantification and the final enrichment analysis in Rstudio software. (**B**) Heatmap of log-CPM values for top 1000 variable genes in each sample. Expression across each gene (row) for all samples (column) is ranged from red for high expression to blue for low expression. (**C**) Dotplot evidencing the lower of expression mitochondrial genes s *CytB*, *ND1* and *lrRNA* in ΔmutDNA samples. (**D**) Dotplot ranking the most 10 activated and suppressed GO biological process (BP) sorted by p value < 0.05 between ctrl and ΔmtDNA, generated by gseGO and dotplot. (**E**) Cnetplot for gene association network between the most 12 differentially expressed GO biological process(GO) sorted by p value > 0.05. Orange remarks highlights biological process of interest for this study.

